# The Abelson kinase and the Nedd4-family E3 ligases co-regulate Notch trafficking to limit signaling

**DOI:** 10.1101/2024.07.12.603272

**Authors:** Julio Miranda-Alban, Nicelio Sanchez-Luege, Fernando M. Valbuena, Chyan Rangel, Ilaria Rebay

## Abstract

Precise output from the conserved Notch signaling pathway governs a plethora of cellular processes and developmental transitions. Unlike other pathways that use a cytoplasmic relay, the Notch cell surface receptor transduces signaling directly to the nucleus, with endocytic trafficking providing critical regulatory nodes. Here we report that the cytoplasmic tyrosine kinase Abelson (Abl) facilitates Notch internalization into late endosomes/multivesicular bodies (LEs), thereby limiting signaling output in both ligand-dependent and -independent contexts. Abl phosphorylates the PPxY motif within Notch, a molecular target for its degradation via Nedd4-family ubiquitin ligases. We show that Su(dx), a family member, mediates the Abl-directed LE regulation of Notch via the PPxY, while another family member, Nedd4Lo, contributes to Notch internalization into LEs through both PPxY-dependent and independent mechanisms. Our findings demonstrate how a network of post-translational modifiers converging at LEs cooperatively modulate Notch signaling to ensure the precision and robustness of its cellular and developmental functions.

## Introduction

Metazoan development relies on spatiotemporal coordination of cellular behaviors. Cells communicate with each other through precisely regulated biochemical signals, with either too little or too much signaling linked to developmental disorders and pathologies^1^. Manipulations of the highly conserved Notch signaling pathway exemplify the sensitivity of developmental events to signal transmission levels^2,3^. Notch orchestrates a diverse array of cellular and developmental processes, including cell fate determination and establishment of tissue boundaries. In some contexts, the *on* versus *off* Notch signaling state induces distinct outcomes, whereas in other cases more modest differences in signaling levels drive specific transitions^4–9^. Because both Notch and its ligands are broadly expressed, this versatility in function requires meticulous spatiotemporal control of the initiation, duration and strength of signaling^10^.

Notch is an integral membrane protein with a ligand-binding extracellular domain (NECD) and a signal transducing intracellular domain (NICD)^11–13^. Canonical activation of Notch signaling^14^ involves cell surface interactions with Delta/Serrate/Lag2-family ligands that trigger endocytosis and proteolytic cleavage of the Notch receptor to release the NICD^15,16^. Notch activation can also occur independent of ligand binding. Unbound Notch receptor is continuously endocytosed^17^ and either recycled back to the cell surface or delivered to the lysosome for degradation^18^. Ligand-independent release of the NICD from specific endosomal compartments is possible during this transport^19–21^. Once cleaved, activated Notch (free NICD) enters the nucleus and forms a protein complex that drives target gene transcription^22–24^. The amount of free NICD quantitatively correlates with the overall transcriptional outcome of signaling^25^ and different dynamics of NICD production can elicit distinct downstream responses^26^.

Multiple factors can influence the production of activated Notch and thereby fine-tune pathway activity. In ligand-induced signaling, the ligand/receptor abundance at cell surfaces (regulated by endocytosis)^21,27,28^, the specific ligand-receptor complex formed^7,26,29,30^, glycosylation of the receptor’s EGF repeats^31–33^ and lipid-ligand interactions^34^ can all affect Notch activation and NICD release dynamics^25^. In ligand-independent signaling, residence of Notch at distinct endocytic/endosomal compartments can affect receptor activation and modify signaling levels^20,35–37^. Therefore, in both ligand-dependent and ligand-independent contexts, precise regulation of Notch endosomal transit must be ensured to modulate NICD release and pathway signaling output.

Endosomal trafficking of integral membrane receptors is controlled by post-translational regulatory mechanisms, most frequently involving ubiquitination and phosphorylation^38,39^. Most insights into the roles of ubiquitination in modulating Notch endocytic flux and signaling stem from studying three specific E3 ubiquitin ligases: Deltex (Dx) and the Nedd4-family members Suppressor of dx (Su(dx)) and Nedd4^40–49^. Genetic, molecular and cell biological approaches have characterized Dx as a positive regulator and Su(dx) and Nedd4 as negative regulators of Notch^40,41,50–59^. Monoubiquitination of the NICD by Dx stabilizes Notch at the limiting membrane of late endosomes/multivesicular bodies (LEs/MVBs)^43,50,51,57–60^, a topology permissive to NICD cleavage and pathway activation^35,50,51,58,60^. In contrast, recognition of the proline-rich PPxY motif in the NICD by the WW domains of Su(dx) or Nedd4 leads to polyubiquitination and downstream degradation of Notch^40,50,51,61,60^. Although unclear for Nedd4^40,41^, the Su(dx)-mediated downregulation of Notch involves internalization of the receptor into the intraluminal vesicles (ILVs) of the LEs/MVBs^50,51,41,58,40,61,60^. This topologically prevents release of cleaved NICD to the cytoplasm and targets Notch for lysosomal degradation^35,50,51,58,60^.

Although receptor trafficking is commonly regulated by both ubiquitination and phosphorylation, phosphorylation-based regulation of Notch endocytic trafficking remains largely unexplored^62^ . The Abelson (Abl) cytoplasmic tyrosine kinase offered an opportunity to address this gap. Abl is best studied as a regulator of actin cytoskeleton and adherens junction dynamics and Abl-Notch genetic interactions have been demonstrated during axon guidance and planar cell polarity establishment^63–69^. However, in these contexts, the underlying mechanism involves Notch-mediated organization of cytoplasmic protein complexes that modulate Abl function, rather than Abl-mediated regulation of Notch signaling. An additional and different role for Abl-Notch interactions was uncovered in the pupal eye disc, where we found that Abl loss impairs clearance of endocytosed Notch^70^. Because reducing the genetic dose of Notch or its transcriptional effector Su(H) suppressed other *abl* mutant phenotypes in the eye, we proposed that Abl-mediated regulation of Notch trafficking might limit signaling. However, the complexity of the *abl* mutant phenotypes in the pupal eye disc^70,71^ made this hypothesis difficult to test.

In this study we used the Drosophila wing vein pattern and S2 cultured cells to investigate the mechanisms by which Abl regulates Notch endocytic trafficking and signaling. We show that loss of Abl results in truncated veins, suggesting excessive Notch activity, and that increasing Abl leads to ectopic vein formation, consistent with insufficient Notch signaling. Subcellularly, loss of Abl results in aberrant accumulation of Notch at the LE membrane, a topology that increases Notch signaling output. Conversely, increased Abl activity drives Notch internalization into the LE lumen and reduces signaling. The LE regulation of Notch by Abl is kinase activity-dependent, and may involve phosphorylation of the NICD’s PPxY, a motif used by Nedd4-family ubiquitin ligases to recognize Notch. Mutation of this motif renders Notch insensitive to regulation by Abl and Su(dx), and partially resistant to a different Nedd4 family member, Nedd4Lo, that can also promote Notch internalization into the LE/MBV lumen. Genetic interaction experiments suggest that the Abl-mediated LE regulation of Notch requires Su(dx), with Nedd4Lo providing parallel PPxY-dependent and -independent inputs. We propose that LEs and the PPxY motif, respectively, provide subcellular and molecular integration points for multiple Notch regulatory inputs. Together or separately, they offer fine-tuning routes applicable to both ligand-dependent and ligand-independent signaling contexts.

## Results

### Loss of *abl* increases Notch signaling and disrupts wing vein pattern

Wing vein patterning is highly sensitive to Notch signaling levels, with too little resulting in excessive vein formation, and too much leading to vein loss^72–74^. The pattern arises during early pupal development when provein cells expressing Delta activate Notch signaling in flanking intervein cells to repress vein fate, ultimately producing the stereotypical vein-intervein pattern by 30-32h after puparium formation (APF)^75,76^. If Abl modulates Notch signaling during wing vein patterning, Abl loss should produce phenotypes characteristic of Notch misregulation.

To test this, we first examined adult wings. Inconveniently, although *abl* null mutants complete pupal development, they die as pharate adults^77^ and, in accordance with our earlier observations^78^, RNAi-mediated Abl knockdown did not produce wing phenotypes, presumably because of Abl protein perdurance. Therefore, we used the deGradFP system^79^ to knockdown Abl protein levels in animals carrying an endogenously GFP-tagged allele^80^ over a null allele (*abl^GFP^/abl*^2^). Adding spatiotemporal control^81^ allowed us to target the knockdown specifically to the posterior compartment of developing pupal wings, thereby bypassing any earlier requirements for Abl function in proliferation or patterning of the larval wing disc^82^. The vein truncations found in 29% of otherwise wildtype-looking Abl knockdown adult wings (Fig. 1A,B; Table1) suggested a disruption in the vein patterning process, and resembled Notch *gain of function* phenotypes. Genetic interactions further implicated Abl as a potential negative regulator within the Notch pathway: reducing *abl* dose, which on its own did not perturb pattern, suppressed the induction of excessive vein material associated with Notch heterozygosity or Su(dx) overexpression and enhanced Dx overexpression-induced vein loss (Table 1).

**Table 1.**
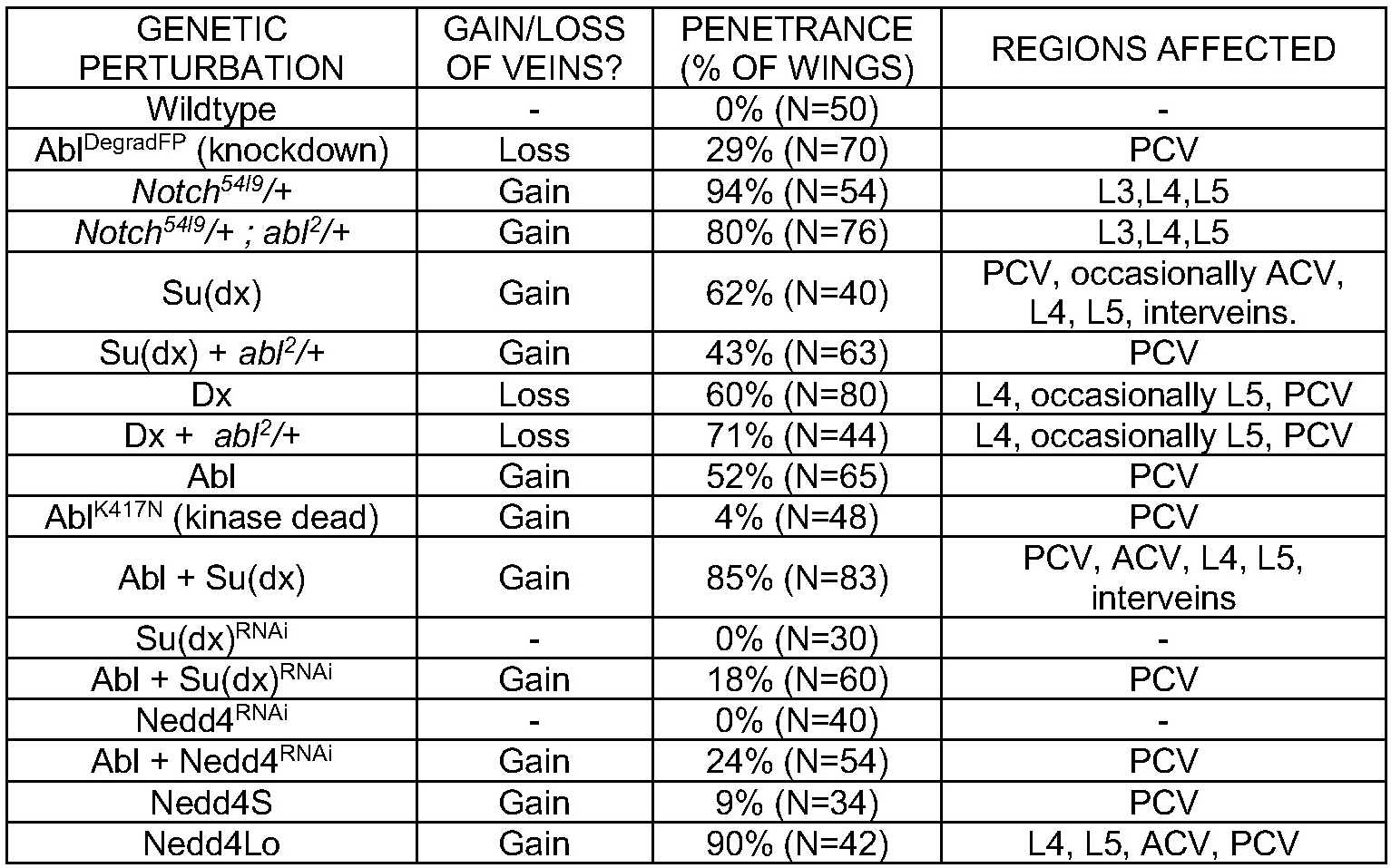
Penetrance of vein pattern phenotypes upon manipulation of Abl and other Notch endosomal trafficking regulators. All genetic perturbations, except the *Notch^54l^*^9^*/+;abl*^2^*/+* interaction, used *engrailed-Gal4* to drive expression of UAS-transgenes in the posterior compartment of developing pupal wings (see methods). Abl knockdown (Abl^DegradFP^) was performed in an *abl^GFP^/abl*^2^ background. ACV/PCV, anterior/posterior cross vein.

**Figure 1.**
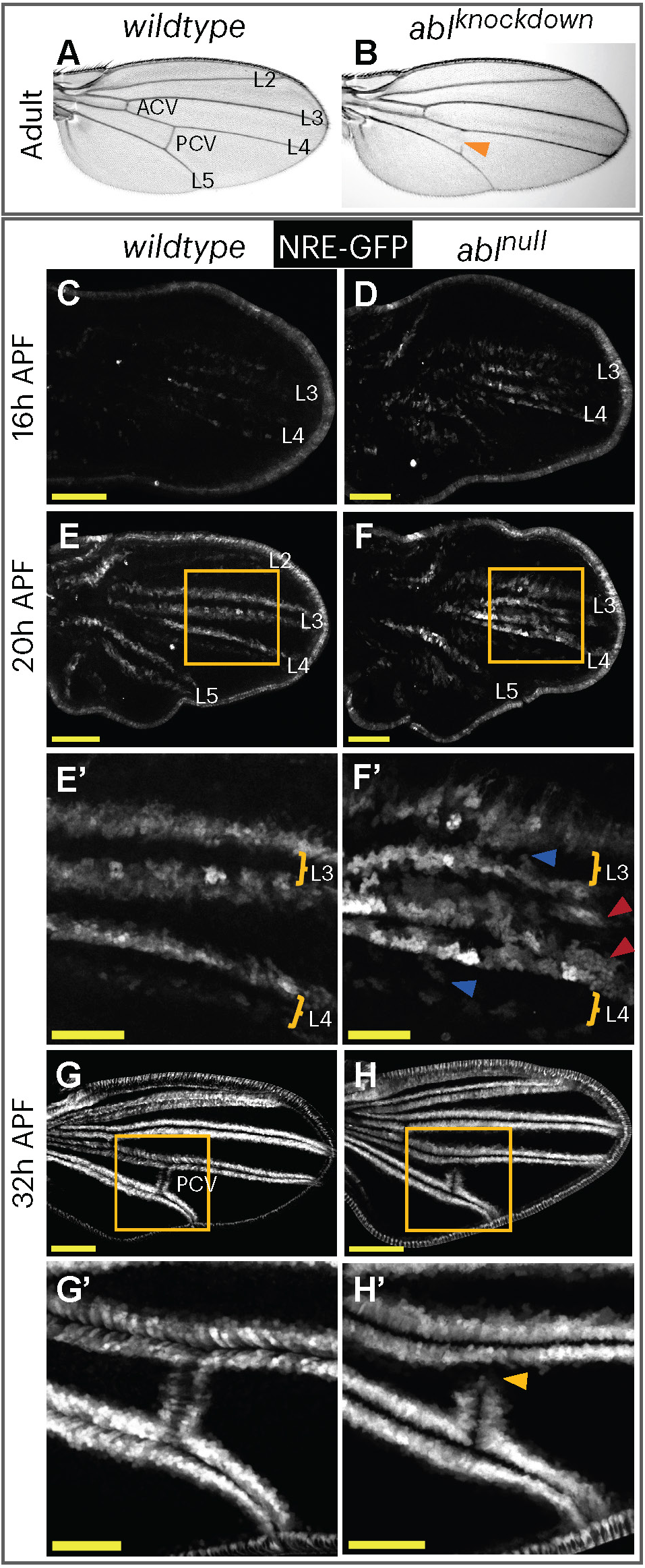
**Abl is required to limit Notch signaling during wing vein patterning.** (A,B) Adult wings. (A) Wildtype, with L2, L3, L4, L5, ACV and PCV veins indicated. (B) *en-Gal4>UAS-deGradFp; abl^GFP^/abl*^2^ (*abl^knockdown^*) adult wings. Orange arrowhead points to a gap in the PCV. (C-H) Maximal projections comparing NRE-GFP reporter expression in wildtype and *abl* null (*abl*^1^*/abl*^2^) pupal wings at 16h (C,D), 20h (E-F) and 32h (G,H) APF. Zoomed insets in (E’,F’) highlight the “noisy” signaling events in L3 and L4 vein (blue arrowheads) and intervein (red arrowheads) cells associated with *abl* knockdown. Zoomed insets in (G’,H’) highlight the loss of NRE-GFP expression in a presumptive PCV gap (yellow arrowhead) that occurs upon loss of *abl*. Scale bars = 100µm (C-H) or 50µm (E’-H’).

Multiple signaling pathways regulate wing vein patterning^83,84^. Therefore, to confirm that the Abl loss-of-function phenotypes resulted from misregulation of Notch, we evaluated the expression of a GFP reporter driven by a Notch Responsive Element (NRE)^85–87^. As previously shown^76,88,89^, in wildtype wings Notch signaling was detected as early as 16h APF in cells flanking the presumptive L3/L4 veins (Fig. 1C) and increased in intensity over time as the pattern refined into two stripes flanking each vein (Figs. 1E, 1G, S1). At 18h APF, zooming into the L3/L4 region (Fig. S1A,A’), small patches of faint reporter expression were detected in the presumptive vein cells (blue arrowheads) and in intervein territory (red arrowheads). This “noisy” signaling activity was resolved by 20hr APF (Fig. 1E,E’). The reporter was also detected at the wing margin at all stages examined, with no obvious change upon *abl* loss (Figs. 1 and S1).

In comparison to wildtype, wings from *abl*^1^*/abl*^2^ null mutants (*abl^null^*) exhibited stronger NRE-GFP signal in the presumptive L3/L4 regions at 16h and 18hr APF (Figs. 1D and S1A-B’). At 18h and 20h APF, the pattern of NRE-GFP appeared “noisier” than in wildtype, with broader-ranged signaling activity in flanking intervein cells and more frequent ectopic signaling events in vein and intervein regions (Figs. 1F,F’, S1A-C). This signaling “noise” was resolved by 24hr APF, after which the pattern and intensity of NRE-GFP reporter expression matched that of comparably staged wildtype wings (Fig. S1D-G). Together these results argue that Abl limits Notch signaling activity during early stages of pupal wing vein patterning.

Starting at 32hr APF, gaps in the NRE-GFP pattern indicative of truncated vein structures were occasionally found in *abl^null^* wings (5%, N=40; Fig. 1G-H’). The frequency of these gaps increased over time (21% at 38h APF, N=78; Fig. S1H-J), consistent with the 29% observed in Abl knockdown adult wings (Table 1). Therefore, endpoint-looking vein pattern defects associated with loss of Abl can be observed as early as the developmental stage when the mature pattern first appears in wildtype (∼32h APF).

While the increased and ectopic NRE-GFP signals detected in early stage *abl* null wings and the final wing vein gaps both supported a Notch gain-of-function interpretation, the loss of reporter expression in flanking intervein cells between 32h-38h APF (Figs. 1H’, S1I,J) at first seemed contradictory. However, consideration of the Notch-Delta negative feedback loop that patterns the vein-intervein boundary^75,76^ offers a logical explanation. According to the model, excessive Notch signaling in presumptive vein cells should abrogate Delta ligand expression and vein fate in those cells, which in turn should reduce ligand-induced Notch signaling in the adjacent cells. If sustained, this should manifest as gaps in NRE-GFP expression and ultimately vein loss. Although in-depth study of Notch-Delta feedback temporal dynamics will be required to confirm this explanation, as predicted by the model, *abl^null^*vein gaps at times had ectopic NRE-GFP reporter expression within the provein region (Fig. 1H’, yellow arrowhead). Altogether, our analysis of Abl loss-of-function phenotypes argues that Abl represses Notch signaling during wing vein patterning.

### Loss of Abl results in endocytic accumulation of Notch

To elucidate the cellular-level consequences of Abl loss that might increase Notch signaling, we compared the subcellular distribution of Notch protein in wildtype versus *abl* null pupal wings. As previously described^76^, in wildtype 32h APF wings, Notch is expressed ubiquitously, with the highest levels of expression (Fig. 2A) and activation (Fig. 1G) in cells immediately flanking the presumptive veins. Subcellularly, Notch appeared enriched at the plasma membrane and in the cytoplasm (Fig. 2A’-A’’), with a punctate distribution that was particularly obvious in the vein regions (arrowheads in Fig. 2A’).

**Figure 2.**
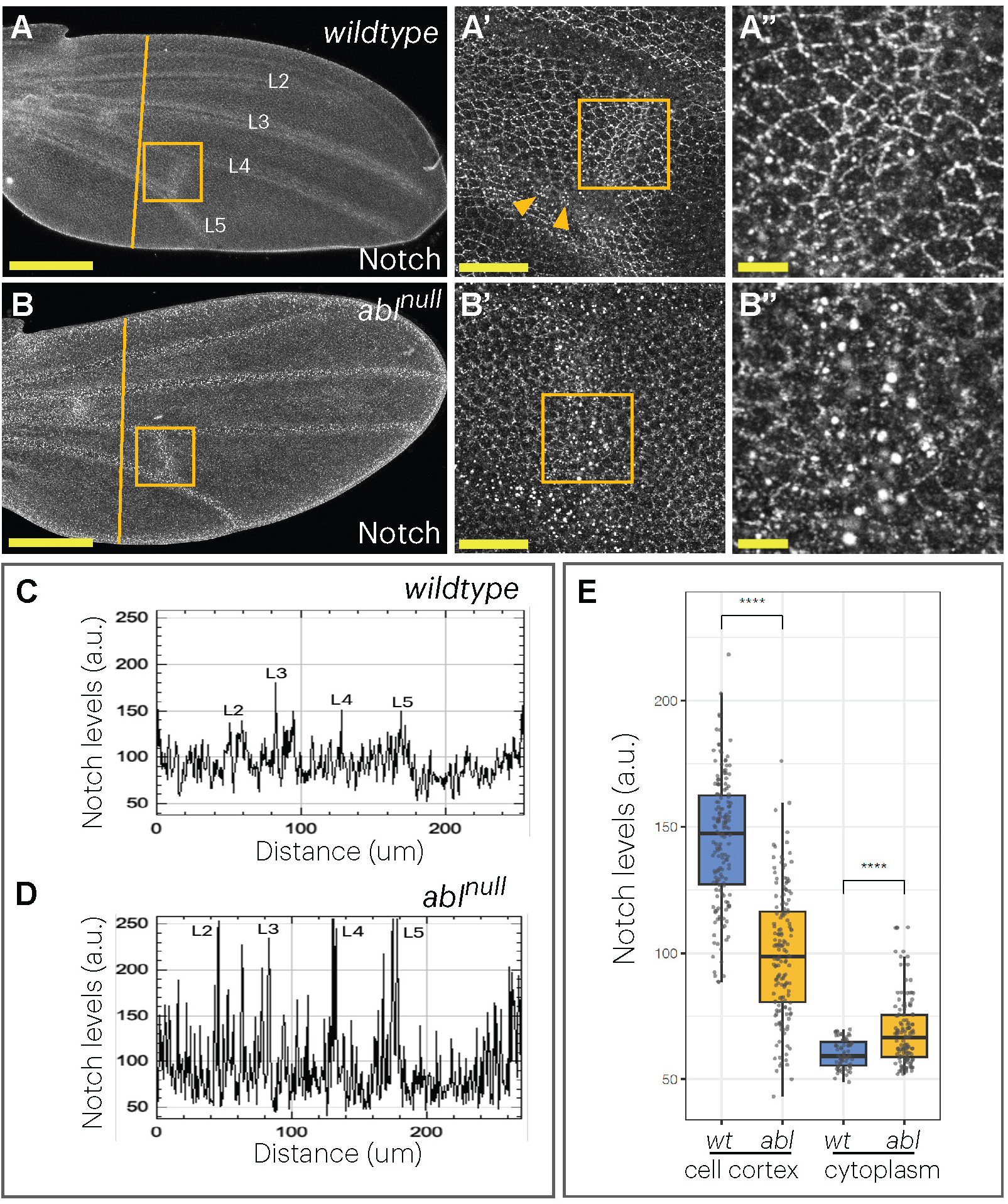
Loss of *abl* increases Notch levels and endocytic accumulation. (A-B’’) Maximal projections showing Notch protein expression detected by indirect immunofluorescence with anti-Notch in 32h APF wildtype and *abl^null^* pupal wings, with zoomed views of the PCV region in (A’,A”,B’,B”). Scale bars = 100µm (A, B), 20µm (A’, B’) or 5µm (A’’, B’’). (C,D) Plots of Notch fluorescent intensity values across the anterior-posterior axis of the wing blade (yellow lines in (A,B)). (E) Quantification of Notch levels at the cell cortex and cytoplasm of cells shown in (A’’) and (B’’) are plotted in (E). Each dot represents the individual measurement of a cell border (N=158 in wt and N=156 in *abl*) or a cell cytoplasm (N=70 in wt and N=80 in *abl*). A t-student means comparison test was performed and p-values <0.05 (*), <0.01 (**), <0.001(***) and <0.0001 (****) are indicated.

In agreement with our prior work^70^, Notch levels appeared elevated in *abl^null^* wings, with the most pronounced increase in the vein regions (Figs. 2B-D). Higher magnification revealed a reduction in membrane-localized Notch and an increase in cytoplasmic Notch (Figs. 2B’’, 2E, S2A-C). The enrichment of Notch in cytoplasmic puncta was particularly striking in the vein regions (Figs. 2B’,B”). Side by side comparison of *abl^null^*and wildtype tissue in somatic mosaics confirmed these differences (Fig S2D,D’).

The localization and turnover of membrane proteins is largely regulated by endocytosis^90,91^, and previous studies have shown that perturbing Notch flux through the endocytic pathway can impact signaling^20,35,37^. Therefore, to assess whether altered patterns of residence in specific endocytic compartments could account for the increased Notch activity observed in *abl^null^* wings, we examined the localization of Notch relative to three endomembrane compartments: upstream endocytic vesicles (Eps15); early endosomes (Hrs); and late endosomes (Rab7)^92–95^. In comparison to wildtype, in *abl^null^*wing cells, Notch enrichment was significantly increased at Eps15^+^ and Rab7^+^ structures, and significantly decreased at Hrs^+^ structures (Fig. 3A-G). We did not detect obvious differences in overall intensity or pattern of these three markers in wildtype versus *abl* null clones (Figs. S2E-G’), nor in the subcellular distribution of E-cadherin (Fig. S2H,H’), a junctional adhesion receptor subject to endocytic regulation in many epithelial tissues^96–99^. This suggests that Abl is not a general endocytic pathway modulator, and that Abl-mediated regulation specifically impacts Notch trafficking.

**Figure 3.**
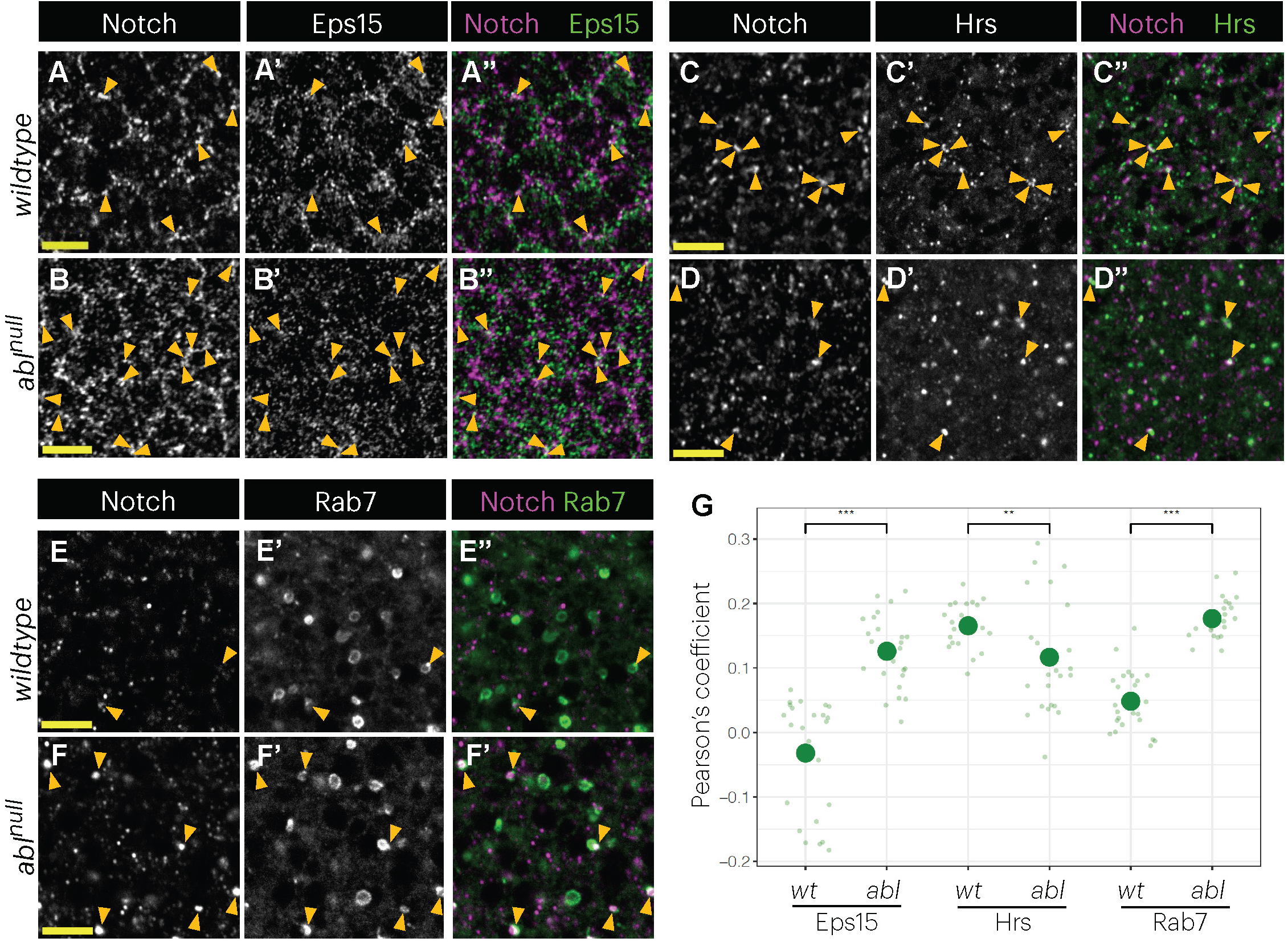
Loss of Abl alters Notch colocalization with endosomal markers. (A-F) Single confocal sections showing Notch and endosomal markers in 32h APF wildtype and *abl^null^* wings. (A,B) anti-Notch and anti-Eps15. (C,D) anti-Notch and anti-Hrs. (E,F) anti-Notch and endogenous Rab7-EYFP. Yellow arrowheads point to examples of Notch-endosomal marker colocalization. Scale bars = 5µm. (G) Pearson’s coefficient-based co-localization analysis. Each dot represents the Pearson’s coefficient from a group of approximately 100 cells in an individual wing. 24 different wings were evaluated per condition. A t-student means comparison test was performed and p-values <0.05 (*), <0.01 (**), <0.001(***) and <0.0001 (****) are indicated.

Prior studies of Notch endocytic regulation offer insight into which of the altered patterns of Notch endocytic residence might underly the gain-of-function wing vein phenotypes associated with Abl loss. While accumulation of Notch in upstream endocytic vesicles has been linked to decreased signaling^35^, accumulation in Hrs+ compartments has been linked to increased signaling^35,100,101^. Therefore, although the respectively increased and decreased localization to Eps15^+^ and Hrs^+^ compartments reflect aspects of the Notch trafficking/turnover defects caused by Abl loss, neither would be predicted to promote Notch activity. In contrast, Rab7-marked late endosomes (LEs) and multivesicular bodies (MVBs) are competent compartments for Notch activation^20,35,37,50,51,58,60^. Therefore, the increased association with Rab7^+^ structures suggested a potential mechanism for Abl-mediated regulation of Notch activity.

### Abl promotes Notch internalization into the late endosome lumen to limit signaling

The specific topology of Notch within LEs determines signaling competence^35,50,51,58,60^. When Notch localizes to the limiting membrane of the organelle, ICD cleavage and cytoplasmic release can activate nuclear signaling. In contrast, internalization of Notch into the LE lumen sequesters the NICD from the cytoplasm and prevents signaling. Therefore, if the increase in Notch localization to Rab7^+^ compartments in *abl^null^*cells activates signaling, Notch should be preferentially enriched at the LE limiting membrane.

Because the small size of wing cells did not offer the resolution needed to distinguish clearly between membrane and luminal domains of Rab7^+^ compartments, we turned to *Drosophila* S2 cells, a cultured cell system the field has used to connect Notch cell biology with pathway activity ^50,87,102^. To validate the system, we first confirmed that Abl limits Notch signaling output. Since S2 cells express Abl endogenously^103^ but do not express Notch^102^, we transiently transfected Notch cDNA with or without Abl dsRNA, and then assessed signaling output using the well-validated NRE>luciferase transcriptional reporter^87,104^. Abl dsRNA led to significantly higher levels of Notch signaling in comparison to transfection with a control dsRNA (Figs. 4G, S3F), indicating that Abl negatively regulates Notch activity in S2 cells.

**Figure 4.**
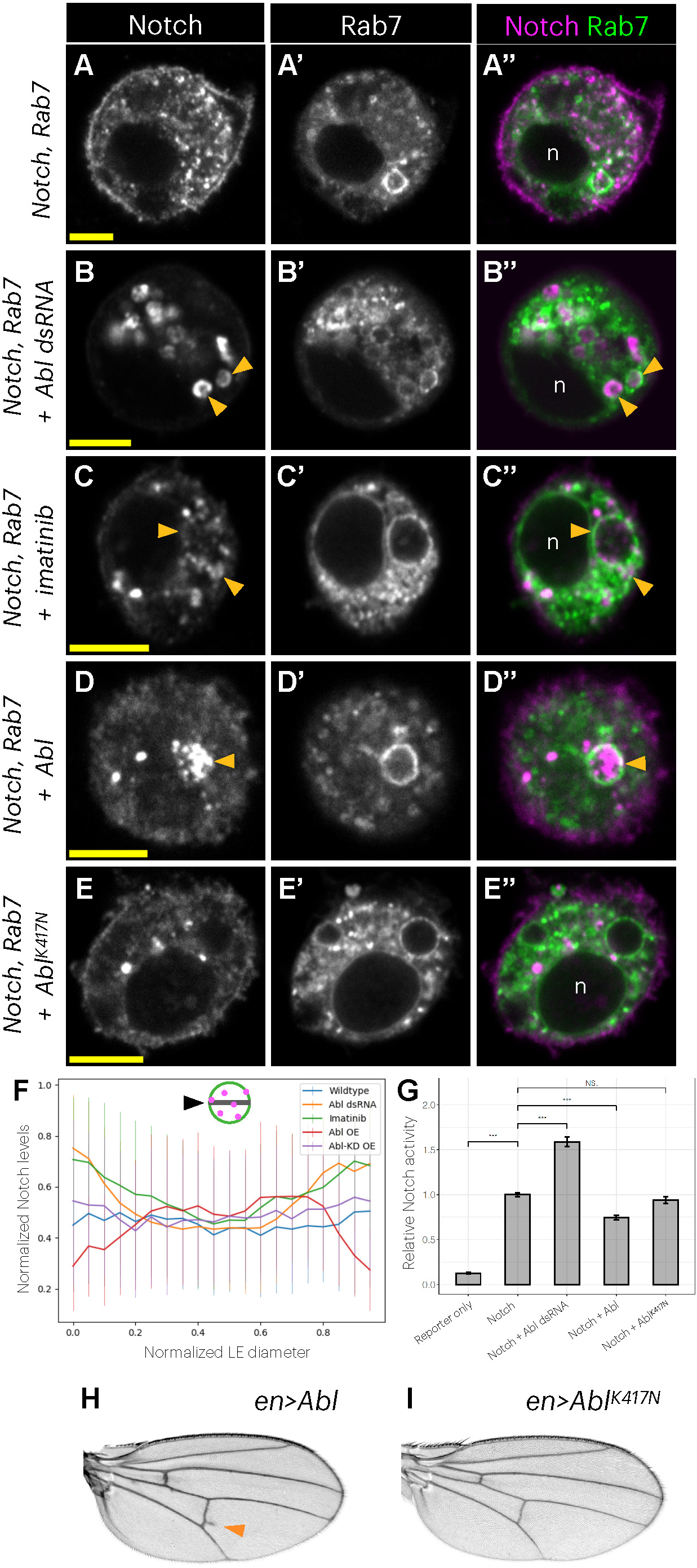
Abl promotes Notch internalization into the late endosome lumen to limit signaling. (A-E) Single confocal sections of representative S2 cells showing Notch (anti-Notch indirect immunofluorescence, magenta) and Rab7-EYFP (green). “n” marks the cell nucleus. Transfections and treatments shown: (A) Notch, (B) Notch + Abl dsRNA, (C) Notch + imatinib mesylate (Gleevec), (D) Notch + Abl, and (E) Notch + Abl^K417N^ (kinase dead). Scale bars = 5µm. (F) Plot showing profiles of Notch levels across late endosome diameters in conditions in (A-E), as depicted by the black arrowhead in cartoon schematic. A 0° cross-sectional line was used to measure Notch fluorescent intensity across LEs. Both Notch levels and LE diameters were normalized. Sample size (# of LEs evaluated): Notch (N=38), Notch + Abl dsRNA (N=32), Notch + imatinib (N=50), Notch + Abl (N=27), Notch + Abl^K417N^ (N=52). (G) Relative Notch activity in manipulations shown in (A-E). A t-student means comparison test was performed and p-values <0.05 (*), <0.01 (**) and <0.001(***) are indicated. (H-I) en>Abl and en>Abl^K417N^ adult wings, respectively. Orange arrowhead in (H) points to an ectopic vein at the PCV.

We next asked if the cellular response to Abl knockdown was comparable to that observed in the wing by examining Notch subcellular distribution in cells expressing Rab7-YFP^93^. Consistent with previous descriptions^50,51,102,105–107^, in cells transfected with Notch alone, and therefore in the presence of endogenous Abl, Notch protein accumulated at the cell cortex and in the cytoplasm, often in small punctate structures (Fig. 4A), some of which were Rab7^+^ (Fig 4A’,A”). Within Rab7^+^ compartments, Notch could be found both at the membrane and in the lumen, with no preferential enrichment in either domain (Fig 4F). Abl knockdown produced a visually striking shift, with reduced cortical Notch and pronounced enrichment in ring-shaped Rab7^+^ cytoplasmic foci (Figs. 4B, S3E). Notch residence at other endosomal compartments was also impacted, with trends similar to those in the wing (Fig. S3A-E, compare to Fig. 3G).

Treatment with imatinib mesylate, a specific inhibitor of Abl kinase activity^108^, similarly shifted Notch subcellular localization from the cell cortex into cytoplasmic foci (Fig. 4C-C”). The predominant enrichment of Notch at the outer membrane of Rab7^+^ LEs was significant for both Abl loss-of-function manipulations (Figs. 4B”,C”,F).

Since internalization of Notch into the luminal space of LEs/MVBs targets the receptor for lysosomal degradation^35,50,51,58,60,55^, Abl might limit Notch signaling by promoting Notch transit into the lumen of these compartments. To test this, we overexpressed Abl in S2 cells and then assessed Notch topology within Rab7^+^ LEs and NRE signaling output. Abl overexpression significantly shifted Notch localization to a predominantly luminal distribution in these compartments (Figs. 4D-D”,F) and reduced signaling output (Fig. 4G). In contrast, overexpression of a well-validated kinase-dead Abl mutant (Abl^K417N^)^78,109,110^ did not promote Notch internalization into the LE lumen or attenuate signaling activity (Figs. 4E-G), further confirming the necessity of Abl’s kinase activity with respect to Notch regulation (Fig. 4C-C”),.

We corroborated these findings in the wing, using the same spatiotemporal control strategy described for the Abl knockdown analysis (Fig 1) to show that overexpression of Abl, but not of kinase-dead Abl^K417N^, induced ectopic veins (Fig. 4H,I and Table 1). Altogether, our results suggest that, in a kinase activity-dependent fashion, Abl promotes Notch relocation from the membrane into the lumen of LEs, thereby negatively regulating Notch signaling by limiting its activation at the LE membrane.

### Abl requires Su(dx) to regulate Notch trafficking and signaling

Targeting of Notch to different LE domains is orchestrated by E3 ubiquitin ligases^40,41,43,50,51,55,57–60^. For example, while Notch is stabilized at the LE membrane by the activity of Dx (Fig. S4A,D), Su(dx) counters this by promoting Notch luminal internalization (Fig. S4B,D). These opposing inputs respectively promote and restrict Notch signaling during wing development^50–56^ (Figs. S4H,I and Table 1).

Given the similarities between Abl and Su(dx) with respect to Notch, we examined genetic interactions between them to probe potential pathway relationships, using signaling output in S2 cells as a quantitative metric. As expected, dsRNA-mediated depletion of Su(dx) increased Notch localization to the membrane of LEs (Figs. 5A,E) and upregulated signaling activity (Fig. 5F). Co-transfection of Abl dsRNA further increased signaling (Figs. 5B,E,F), suggesting a cooperative relationship between Abl and Su(dx) in the LE regulation of Notch. Supporting this, co-overexpression of Abl and Su(dx) led to a greater reduction in signaling than with either Abl or Su(dx) alone, and to the expected Notch LE luminal enrichment (Figs. S4C-E). Such functional synergy could result from Abl and Su(dx) either working together in the same pathway or providing parallel inputs that converge on Notch.

**Figure 5.**
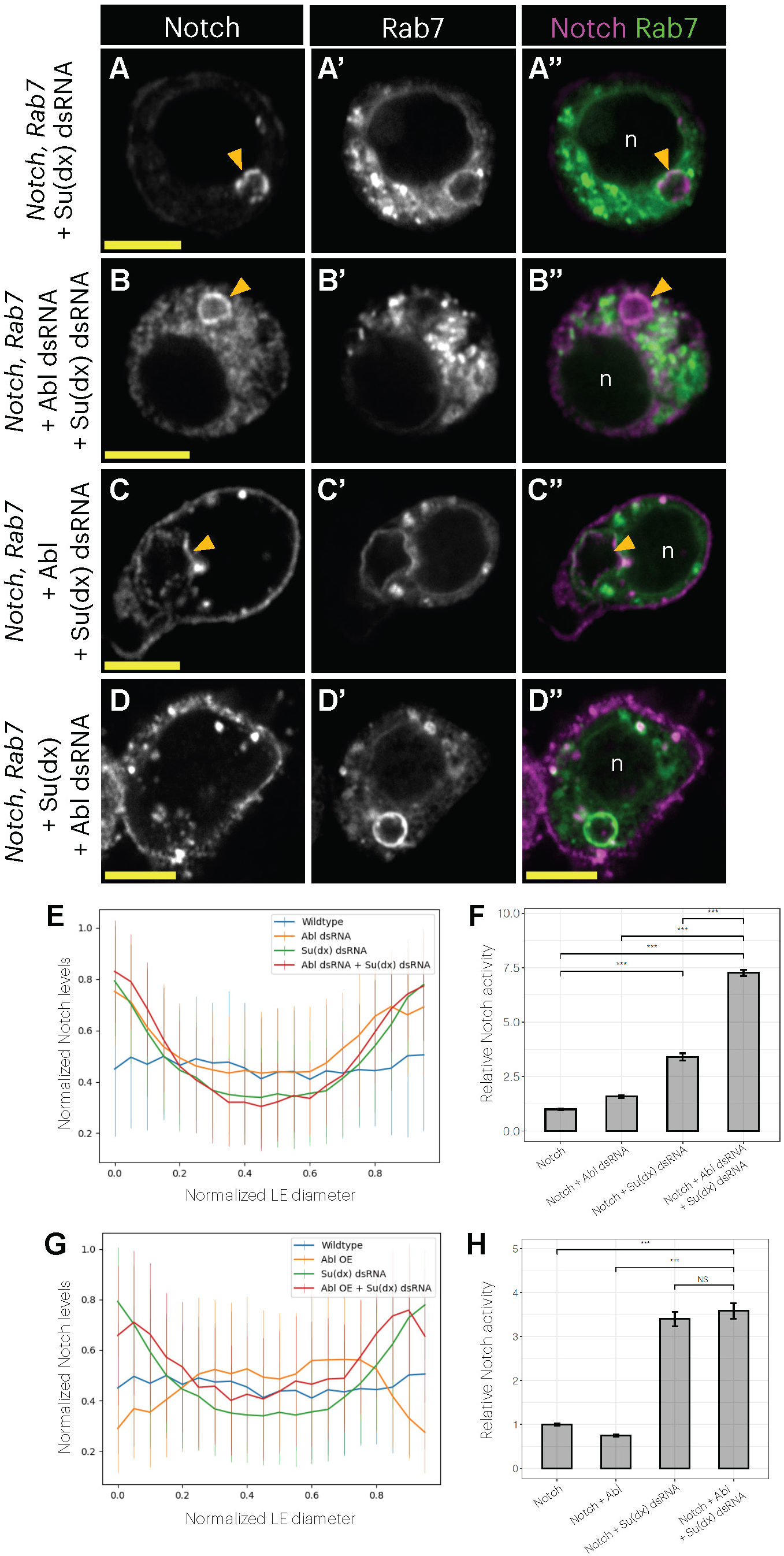
Abl requires Su(dx) to internalize Notch into late endosomes and attenuate signaling. (A-D) Single confocal sections of representative S2 cells showing Notch (indirect immunofluorescence) and Rab7-EYFP . “n” marks the cell nucleus. Transfections shown: (A) Notch + Su(dx) dsRNA, (B) Notch + Abl dsRNA + Su(dx) dsRNA, (C) Notch + Abl + Su(dx) dsRNA, and (D) Notch + Su(dx) + Abl dsRNA. Scale bars = 5µm. (E,G) Plot showing profiles of Notch levels across late endosome diameters in the indicated manipulations. Sample size (# of LEs evaluated): Notch (N=38), Notch + Abl dsRNA (N=32), Notch + Su(dx) dsRNA (N=68), Notch + Abl dsRNA + Su(dx) dsRNA (N=53), Notch + Abl (N=27), Notch + Abl + Su(dx) dsRNA (N=27). (F,H) Relative Notch activity in the indicated manipulations. A t-student means comparison test was performed and p-values <0.05 (*), <0.01 (**) and <0.001(***) are indicated.

To assess the epistatic relationship, we tested whether Su(dx) is required for Abl-mediated regulation of Notch, or vice versa. Depletion of Su(dx) completely masked the effects of Abl overexpression, producing an enrichment of Notch at LE membranes and an increased signaling output comparable to that produced by Su(dx) dsRNA alone (Figs. 5C,G,H). The epistatic relationship suggested that Abl-mediated regulation of Notch is dependent on Su(dx). Performing the experiment in the other direction, depletion of Abl in Su(dx)-overexpressing cells restored the balance of Notch LE luminal versus membrane localization and signaling output to that typical of cells transfected with Notch alone (Figs. 5D, S4F,G). The simplest interpretation of this mutual suppression is that the activity of Su(dx) on Notch is not mediated by Abl, consistent with Abl working upstream of Su(dx).

Suggesting a comparable Abl-Su(dx) synergy and epistatic relationship *in vivo*, the penetrance of ectopic veins resulting from co-overexpression of Abl and Su(dx) exceeded that of each individual manipulation (Table 1). Moreover, while Su(dx) RNAi did not modify the vein pattern in an otherwise wildtype background (Table 1), it substantially suppressed the ectopic vein formation induced by Abl overexpression (Table 1). Altogether, our results suggest that Abl and Su(dx) cooperatively regulate Notch trafficking and signaling, with Abl function dependent on Su(dx).

### The NICD’s PPxY motif integrates Abl, Su(dx) and Nedd4Lo-mediated regulation of Notch trafficking and signaling

Given the necessity of Abl’s kinase activity in the regulation of Notch trafficking and signaling (Figs. 4C,E,F) and the presence of a tyrosine residue, Y2328, in the PPxY motif within the NICD that is recognized by Nedd4-family ligases^40,41,61^, we hypothesized that this motif might integrate Abl and Nedd4-family regulatory inputs. Using an established *in vitro* kinase assay^78^, we found that Abl can phosphorylate GST-NICD but not GST alone (Fig. 6A). Mutation of the Y2328 tyrosine reduced phosphorylation of the GST-NICD^Y2328F^ substrate (Fig. 6A), indicating that the Notch PPxY tyrosine can be a direct target of Abl activity. The residual phosphorylation signal in GST-NICD^Y2328F^ was expected, as previous work has shown that Abl can phosphorylate other NICD tyrosine residues^68^.

**Figure 6.**
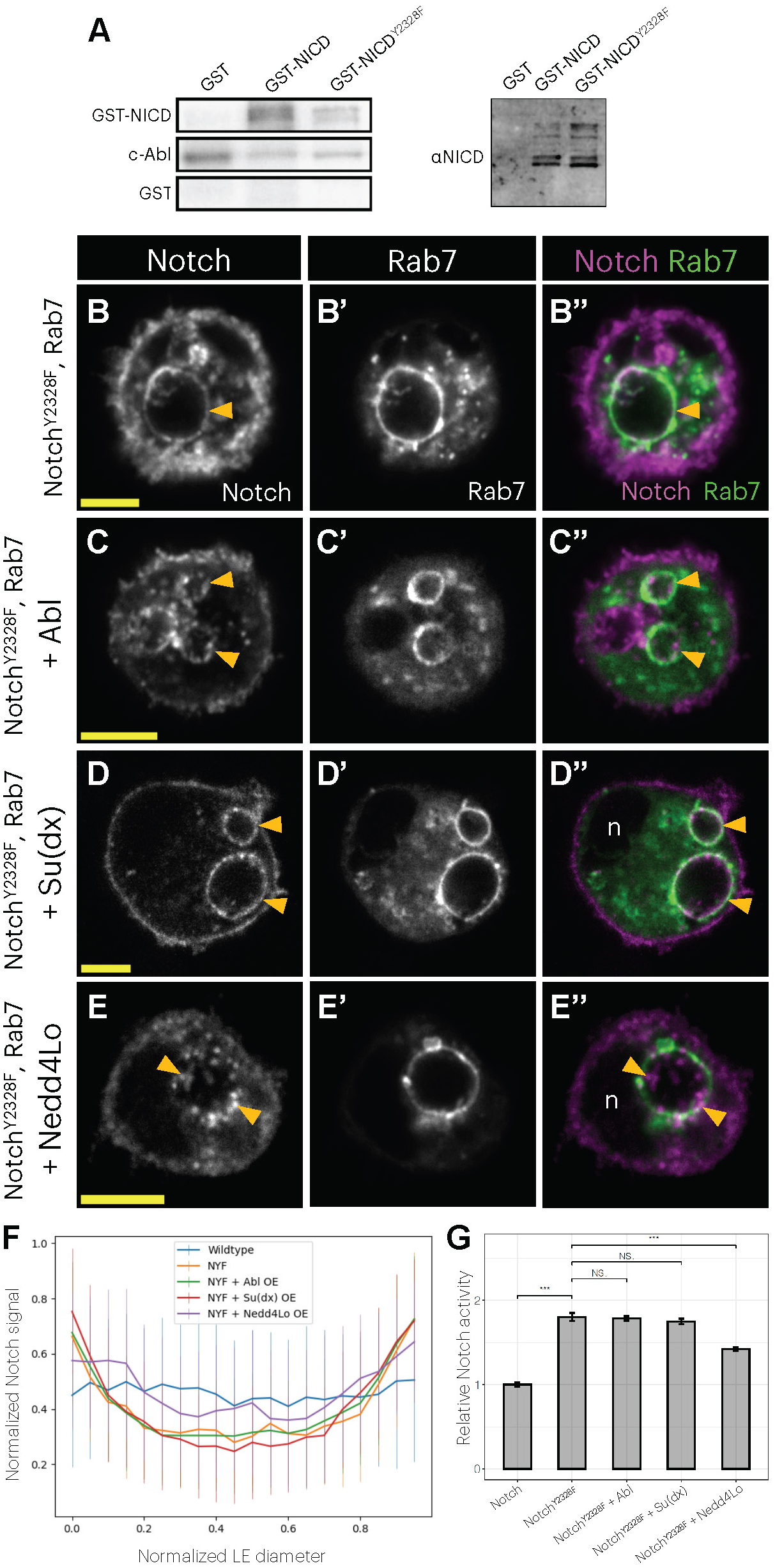
The NICD’s PPxY tyrosine integrates Abl, Su(dx) and Nedd4Lo-mediated regulation of Notch trafficking and signaling at LEs. (A) Left panel: *in vitro* kinase assay showing that recombinant c-Abl can phosphorylate the NICD, with some activity targeted to the Y2328 residue. Right panel: Western blot showing GST-NICD loading in samples evaluated on left panel. (B-E) Single confocal sections of representative S2 cells showing Notch (indirect immunofluorescence) and Rab7-EYFP . “n” marks the cell nucleus. Transfections shown: (B) Notch^Y2328F^, (C) Notch^Y2328F^ + Abl, (D) Notch^Y2328F^ + Su(dx), and (E) Notch^Y2328F^ + Nedd4Lo. Scale bars = 5µm. (F) Notch signal profiles across late endosome diameters in manipulations in (B-E). Sample size (# of LEs evaluated): Notch (N=38), Notch^Y2328F^ (N=43), Notch^Y2328F^ + Abl (N=73), Notch^Y2328F^ + Su(dx) (N=38), Notch^Y2328F^ + Nedd4Lo (N=52). (G) Relative Notch activity in manipulations in (B-E). A t-student means comparison test was performed and p-values <0.05 (*), <0.01 (**) and <0.001(***) are indicated.

To test the functional significance of the PPxY tyrosine, we introduced a tyrosine to phenylalanine substitution to generate Notch^Y2328F^. When transiently transfected into S2 cells, the enrichment of Notch^Y2328F^ at the limiting membrane of LEs (Fig. 6B) was enhanced relative to wildtype Notch (Fig. 6F). Consistent with its enrichment at the LE membrane, signaling output from Notch^Y2328F^ was increased relative to wildtype Notch (Fig. 6G). If the PPxY tyrosine provides a molecular target for Abl-mediated antagonism of Notch signaling, then Notch^Y2328F^ localization and activity should be insensitive to Abl overexpression. As predicted, neither the increased localization of Notch^Y2328^ to LE membranes nor its elevated signaling output were modified by co-transfection of Abl (Figs. 6C,F,G). Notch^Y2328F^ localization and activity was also insensitive to Su(dx) overexpression (Figs. 6D,F,G), arguing that the Notch Y2328 is required for both the Abl- and Su(dx)-mediated LE internalization and signaling modulation.

In addition to mediating regulation by Abl and Su(dx), the NICD PPxY motif has also been implicated in Nedd4-mediated ubiquitination and degradation of Notch ^40,41^. Nedd4 encodes two major isoforms, Nedd4short (Nedd4S) and Nedd4long (Nedd4Lo), which have been suggested to have distinct genetic requirements and cell biological functions^111–113^ . We were therefore curious whether Nedd4 might also participate in Notch LE regulation, perhaps with different isoform specificity. To assess this, we examined the effects of Nedd4 depletion or overexpression on Notch LE localization and signaling in S2 cells. Although neither Nedd4 dsRNA nor expression of the short isoform (Nedd4S) altered Notch signaling (Fig S3F), expression of the long Nedd4 isoform (Nedd4Lo) significantly decreased signaling (Fig. S5D) and enriched Notch localization to LE lumens (Figs. S5A-C). Consistent with the isoform-specificity uncovered in S2 cells, overexpression of Nedd4Lo strongly induced excessive vein formation in 90% of wings, a phenotype consistent with reduced Notch activity, while Nedd4S only rarely perturbed pattern (Figs. S5E,F; Table 1). Nedd4 knockdown also suppressed the ectopic vein formation induced by Abl overexpression (Table 1), suggesting that, like Su(dx), it may act downstream of Abl.

Returning to the S2 cells, we asked if the PPxY motif was necessary for Nedd4Lo-mediated regulation of Notch LE localization and signaling. In contrast to our finding that Notch^Y2328F^ was resistant to regulation by both Su(dx) and Abl, Nedd4Lo transfection led to a significant (∼20%) reduction in Notch^Y2328F^ signaling (Fig. 6G). Consistent with the reduced signaling, LE luminal Notch^Y2328F^ subcellular localization was readily detected upon Nedd4Lo transfection. Given that Nedd4Lo reduced wildtype Notch signaling by ∼40% (Fig. S5D), the ∼20% reduction in Notch^Y2328F^ signaling (Fig. 5G) suggests that Nedd4Lo regulates Notch via both PPxY-dependent and independent mechanisms. Together with the previous report of a partial requirement for the PPxY in the Nedd4-mediated ubiquitination of Notch^40^, our result offers a logical mechanistic connection between Notch ubiquitination, subcellular localization and signaling. Altogether, our results highlight LEs and the PPxY motif as versatile regulatory hubs that can integrate multiple inputs to finely modulate Notch signaling output.

## Discussion

Endocytic internalization and trafficking of transmembrane signaling receptors play key roles in modulating pathway output^114–117^. Given the Notch pathway’s distinctive lack of a cytoplasmic signaling cascade, regulation of receptor passage through endosomal compartments provides each cell, whether ligand-stimulated or not, with autonomous control over Notch levels and pathway activity^19,20,37,44,47,49,118^. In this study we identify the cytoplasmic tyrosine kinase Abelson as a novel regulator of Notch trafficking and signaling. Our findings reveal a kinase-dependent role for Abl in promoting the clearance of endocytosed Notch and establish an unanticipated connection between Abl and a regulatory network of ubiquitin ligases that controls the topology, accessibility and enrichment of Notch at LEs to modulate signaling output. We propose that LEs provide dynamic subcellular hubs, capable of integrating multiple regulatory inputs to finely tune Notch activity to levels optimal for specific cellular and developmental contexts.

Previous work has demonstrated the key role of E3 ubiquitin ligases in controlling Notch dynamics at LEs^40,41,43,50,51,58,60^, but the molecular events that control their activity have remained poorly understood^119^. At the molecular level, Nedd4-family proteins use their WW domains to recognize and bind PPxY motifs within their substrates^40,61,120–123^, an interaction involved in forming complexes such as Su(dx)/Nedd4-Notch in *Drosophila* and the analogous human WWP2-NOTCH3^40,61,123^. The connections between the ubiquitin ligases network and the Abl kinase uncovered in our study mark the Notch PPxY motif as an important molecular integration point for WW domain-based and tyrosine phosphorylation inputs that modulate Notch subcellular localization and signaling. Although we tried to generate an endogenous *Notch^Y^*^2328^*^F^*allele via CRISPR/Cas9, the recovered heterozygous females were too sick and infertile to establish a genetic line. However, those flies exhibited a marked reduction in small thoracic bristle density^124^, a phenotype suggestive of excessive Notch activity^125,126^. Alternative genetic strategies will therefore be needed to explore how the Notch PPxY motif integrates kinase and ubiquitin ligase inputs to regulate trafficking and signaling in developing tissues.

While protein structure disruptions could underlie the insensitivity of Notch^Y2328F^ to LE regulators, we favor a model in which the phosphorylation state of the Notch PPxY motif influences its interaction with and regulation by Nedd4-family members. Prior work has shown that PPxY phosphorylation can either enhance or reduce a particular interaction^120,127,128^. For example, phosphorylation of the c-Jun PPxY motif in T-cells prevents Nedd4-family ligase binding, thereby protecting the transcription factor from degradation^128^. c-Abl was implicated as the kinase, which combined with our findings, hints at a role that might extend to diverse target proteins and biological contexts.

In contrast to the c-Abl/c-Jun relationship, our results suggest Abl-mediated phosphorylation of the Notch PPxY motif potentiates regulation by Nedd4-family ligases. By creating a more favorable interaction platform for Su(dx) or Nedd4Lo recruitment, phosphorylation could increase the probability of ubiquitination patterns that drive Notch into signaling-incompetent endosomal domains. Given that Su(dx) recognizes Notch using a unique WW4 domain absent in other Nedd4-family members^121,122^, Su(dx) and Nedd4Lo might also have distinct PPxY motif phosphorylation preferences or affinities, enabling them to compete and collaborate in different ways. Therefore, in some situations, the activity of Abl on Notch might rely primarily on Su(dx) or on Nedd4Lo. In others, for example the pupal wing epithelium, where RNAi knockdown of either Su(dx) or Nedd4 suppressed the ectopic vein phenotype associated with Abl overexpression (Table 1), contributions from multiple Nedd4-family ligases may be required.

Beyond mediating Abl-Nedd4 ligases interactions, the Notch PPxY motif could provide a molecular hub whose phosphorylated versus unphosphorylated states are read and acted upon by additional regulators. These could include tyrosine phosphatases, phosphotyrosine binding proteins, WW domain containing proteins, among others. By organizing diverse molecular complexes with different regulatory and signaling capabilities, these interactions could provide highly dynamic control of Notch activity. In addition, because Notch-Abl interactions have also been shown to influence Abl activity^68,69^, the possibility of multiple layers of feedback regulation offers essentially limitless opportunities for each cell to fine-tune its Notch levels, subcellular localization and signaling output.

The NICD PPxY motif need not be the only target of Abl. For example, tyrosine phosphorylation of Nedd4-family proteins has been shown to relieve autoinhibitory mechanisms^129–131^. Therefore, Abl could facilitate the LE internalization of Notch by unlocking or enhancing the regulatory potential of Nedd4 ligases in more than one way. While the molecular events that mediate recognition of Nedd4 ligases by tyrosine kinases have not been elucidated, the latter typically use their Src Homology (SH) domains to bind other proteins^132^. Such interactions could facilitate targeting and tyrosine phosphorylation of NEDD4-1 by c-Src^131^, for example. Akin to Src, Abl possesses SH2 and SH3 domains^133^ and, in particular, SH3 domains have been shown to bind proline-rich motifs in Nedd4-family ligases^112,134,135^. In Drosophila, Nedd4Lo differs from Nedd4S in the presence of extra proline-rich regions in both its N-terminal and Mid domains^112^. In the case of the Mid domain, the proline-rich region separates the WW1 from the WW2 and WW3 domains^111,112^, an organization similar to the separation of the Su(dx) WW3 and WW4 domains by a proline-rich region^122^. Hence, the unique molecular structure of certain Nedd4 family members, even at the isoform level, could nucleate specific complexes that interact with Notch in both PPxY-dependent and independent manners.

In conclusion, we propose that the spatiotemporal patterning of Notch post-translational modifications offers a versatile regulatory strategy to modulate endosomal trafficking and thereby tune Notch signaling levels to the particular cellular or developmental transition. In the developing wing, the enrichment of Abl and Su(dx) in cells flanking proveins^55,136^ suggests that tight regulation of LE trafficking may be critical to attenuating Notch signaling in cells that are exposed to high concentration of ligand. On the other hand, the ubiquitous Notch trafficking defects observed in *abl^null^*mutant wings, even outside of the vein areas (Fig. S2A,B), imply that cells can use the endocytic pathway to modulate Notch activation even in the absence of ligand stimulus. LE regulation can thus provide a versatile system to adjust Notch signaling output in both ligand induced and independent contexts. As dysregulated Notch activity is a key feature in oncogenic disorders linked to dysfunction of Abl, Dx, Su(dx) and Nedd4^137–141^, our study highlights cellular processes and molecular interactions that could be potential targets for therapeutic interventions.

## Materials and Methods

### Drosophila genetics

All flies were raised on standard *Drosophila* food at 25°C unless otherwise noted. The following lines were used: *abl*^1^ ^77^, *abl*^2^ ^77^, *abl-GFP*^80^, NRE-EGFP (BSDC 30727), *rab7-myc-EYFP*^93^, en>Gal4, Gal80ts, UAS-deGradFp^79^, UAS-Abl-GFP^110^, UAS-Abl^K417N^-GFP^110^, UAS-Flag-Dx (a gift from Spyros Artavanis-Tsakonas), UAS-Su(dx) (BSDC 15664), UAS-Nedd4Short-Flag^111^, UAS-Nedd4Long-Flag^111^, UAS-Su(dx) RNAi (BSDC 67012), and UAS-Nedd4 RNAi (BSDC 34741)). For UAS-driven genetic manipulations, crosses were established at 18°C and individual animals were transferred to 25°C at 0h APF. They were either dissected at 32h APF or allowed to grow to adulthood. To generate heatshock-driven *abl* clones in wing tissue, *y, w, hs>Flp;; ubi-GFP, FRT80B/TM6, Tb* females crossed to *abl*^2^*, FRT80B/TM6, Tb* males. Parents laid eggs overnight, offspring were allowed to develop for 32h before heatshock treatment at 37°C for 45min, and heatshocked animals were transferred back to 25°C until dissection.

### Immunostaining of fly tissue, antibodies and co-localization analysis

16-40h APF wing dissections: pupae were decapitated, fixed in 4% paraformaldehyde (PFA) in PBS for 12-24h at 4°C, washed 3x and then dissected in PBS. Dissected wings were immediately blocked in 1% normal goat serum in PBT for 30min at room temperature or overnight at 4°C, incubated overnight at 4°C with primary antibodies in PBT, washed 3x in PBT, incubated overnight with secondary antibodies at 4°C, washed 3x in PBT, mounted in n-propyl gallate mounting medium and imaged on a Zeiss LSM 880 confocal microscope. The Airyscan function was used for Notch-Rab7 colocalization experiments. GFP-tagged proteins were imaged using the endogenous fluorescent signal. Antibodies: mouse anti-NECD (1:100, Developmental Studies Hybridoma Bank (DHSB) C458.2H), mouse anti-Delta (1:500, DHSB C594.9C), guinea pig anti-Eps15 and anti-Hrs (1:500, gifts from Hugo Bellen), goat anti-mouse Cy3, Alexa 488 and Cy3 (1:2000, Jackson ImmunoResearch), goat anti-guinea pig Cy3 and Alexa488 (1:2000, Jackson Immunoresearch), and goat anti-rabbit Cy5 (1:2000, Jackson Immunoresearch). Co-localization analysis between Notch and endosomal markers was performed by calculating the Pearson’s correlation coefficient between Notch and endosome channels using the COLOC2 plug-in of Image J/Fiji. Regions of approximately 100 cells were evaluated per wing, and 24 different wings were evaluated per genotype and marker. Profiles of Notch distribution across LE diameters were obtained from all LEs>0.8um (N=30-70) for each condition, and normalized to allow comparisons between experiments.

### S2 cell transfection and dsRNA/imatinib treatments

S2 cells were cultured in Schneider’s insect medium (Sigma catalog S9895-10X1L) supplemented with Penicillin and Streptomycin at 25°C. For all assays, 2.25 x 10^6^ cells were transfected with dimethyldioctadecylammonium bromide (DDAB, Sigma D2779-10G), pMT expression plasmids, dsRNA, and pMT vector to equalize the total amount of nucleic acid transfected, up to 1ug, Transfection was conducted for 30min, followed by incubation for 24h at 25°C, before induction with 0.7mM CuSO4 for 24h at 25°C.

Plasmid and dsRNA transfection amounts: pMT-Notch (150 ng), pMT-Notch^Y2328F^ (150 ng), pMT-Abl-Myc (250ng), pMT-Su(dx)-Flag (150 ng), pMT-Dx-Flag (150ng), pMT-Flag-Nedd4S (150ng), pMT-Flag-Nedd4Lo (150ng), Abl dsRNA (350ng), Su(dx) dsRNA (250ng), Nedd4 dsRNA (250ng), LacZ dsRNA (250-350ng), pMT-EYFP-Rab7 (250ng, a gift from Martin Baron) and pMT vector. Transcription assays also included 2 ng of NRE-Luciferase (Firefly) reporter (a gift from Sarah Bray), and 20 ng of actin-Renilla normalizing control. For imatinib mesylate treatment, imatinib mesylate solution in water was added to transfected cells at a final concentration of 50-100 µM at the time of induction.

For dsRNA preparation, target sequences from pMT-Abl-Myc^78^, pMT-Su(dx)-Flag (a gift from Martin Baron) and pcDNA3-Flag-Nedd4Lo (a gift from Daniela Rotine) were PCR-amplified using the following primers: 5’-TAATACGACTCACTATAGGGAGAAAAATTTGTTTGGCCTTTTCAA-3’ and 5’-TAATACGACTCACTATAGGGAGAACGGTCATCCTATATCTTTT-3’ for Abl dsRNA^142^, 5’-TAATACGACTCACTATAGGGAGATACATCACCCTCATGACG-3’ and 5’-TAATACGACTCACTATAGGGAGATCATTCCTGGCAGAAGCC-3’ for Su(dx) dsRNA^143^, 5’-TAATACGACTCACTATAGGGAGAGTCACAATAGTATAGAGGACAA-3’ and 5’-TAATACGACTCACTATAGGGAGAGGTCGTTCTGCAAACTGGTC-3’ for Nedd4 dsRNA 1^144^, and 5’-TAATACGACTCACTATAGGGAGACGACTTGAAGAGCAGCAACA-3’ and 5’-TAATACGACTCACTATAGGGAGATAGGTTCGTGTGGTTCACCA-3’ for Nedd4 dsRNA 2.

Transcription of amplified sequences was done using the Ambion MEGAscript^TM^ kit (ThermoFisher Scientific) and subsequent purification was done via standard ethanol precipitation.

### S2 cell-based subcellular localization and transcription/luciferase assays

For subcellular localization assays, cells were settled on poly-L lysine coated slides for 1h at 25°C, fixed for 15 minutes in 4% PFA in PBS, washed 3x in PBS, incubated for 1h at room temperature with primary antibodies in PBT+1% normal goat serum, washed 3x in PBS, incubated with secondary antibodies in PBT for 2h at room temperature, washed 3x in PBS, mounted in n-propyl gallate mounting medium, and imaged on a Zeiss LSM 880 confocal microscope. The same antibodies used for fly tissue were used for S2 cell staining and a minimum of 20 cells were imaged and considered for further examination. Co-localization analysis between Notch and endosomal markers was done as described above. Profiles of Notch distribution across LE diameters were obtained from all LEs>0.8um (N=25-75) for each condition. Both the Notch signal and the LE diameter were normalized to a 0-1 scale to allow comparisons between experiments. Custom python script available upon request.

For transcription assays, each biological replicate in our plots represents a separate transfection. Cells were lysed in 100mM Potassium Phosphate, 0.5% NP-40, 1 mM DTT, pH7.8 for 1h on ice. Lysates were loaded in triplicate into an Autolumat Plus LB 953 luminometer.

Firefly and Renilla luciferases were activated in separate reactions using Firefly buffer (10mM Mg Acetate, 100mM Tris Acetate, 1mM EDTA, 4.5mM ATP, and 77uM D-luciferin, pH 7.8) and Renilla buffer (25mM sodium pyrophosphate, 10mM Na Acetate, 15mM EDTA, 500mM Na2SO4 500mM NaCl, 4mM coelenterazine, pH 5.0), respectively. The ratio of Firefly RLU to Renilla RLU was averaged across technical replicates, and activation values were normalized to Notch activity.

### Molecular cloning of pGEX-NICD, pGEX-NICD^Y2328F^, and pMT-Notch^Y2328F^

pMT-NotchY2328F was made by subcloning the XhoI-XbaI NICD fragment from pMT-Notch into pBluescript. The Y2328F point mutation was introduced using Quickchange PCR using the following primers: 5’-AAGCAGCCGCCGAGCTTTGAGGATTGCATCAAG-3’; 5’-CTTGATGCAATCCTCAAAGCTCGGCGGCTGCTT-3’. The mutated XhoI-XbaI fragment, confirmed by sequencing, was cloned back into pMT-Notch to create pMT-Notch^Y2328F^. Wildtype and Y2328F XhoI-NotI NICD fragments were subcloned into SalI-NotI digested pGEX-4T-2 plasmid to create the bacterial expression constructs.

### In vitro kinase assay

GST-fusion proteins were purified from BL21 *E. coli* cells as described in Rebay and Fehon, 2009 . Kinase assays were performed as described in Xiong et al., 2009^78^. The relative intensity of full-length wildtype and Y2328F GST-NICD bands, as detected on western blots with mouse anti-NICD (1:1000) antibody, was used to estimate comparable amounts of substrate for the reactions Samples were run on a 10% polyacrylamide gel, transferred to a PVDF membrane using standard methods and exposed on a Storm phosphoimager prior to immunoblot analysis to confirm protein loading.

## Acknowledgements

We would like to thank Spyros Artavanis-Tsakonas, Wei Du and Hitoshi Matakatsu for fly strains, Sarah Bray, Martin Baron and Daniela Rotin for S2 cell expression plasmids, Hugo Bellen and Rick Fehon for antibodies, and Lucy Godley for imatinib mesylate. We also thank Hideyoshi Shimizu and Hitoshi Matakatsu for advice on pupal wing dissection, Allison Zajac, Jacob Decker and Jose Velarde for advice on co-localization analysis, Xiao Sun and Christine Cao for experimental assistance, and Rick Fehon for helpful comments on the manuscript. In addition, we thank all members of the Rebay Lab, Rick Fehon, Chip Ferguson, Sally Horne-Badovinac, Aaron Turkewitz and Paschalis Kratsios for helpful discussions.

## Supplementary Figure Legends

**Figure S1.**
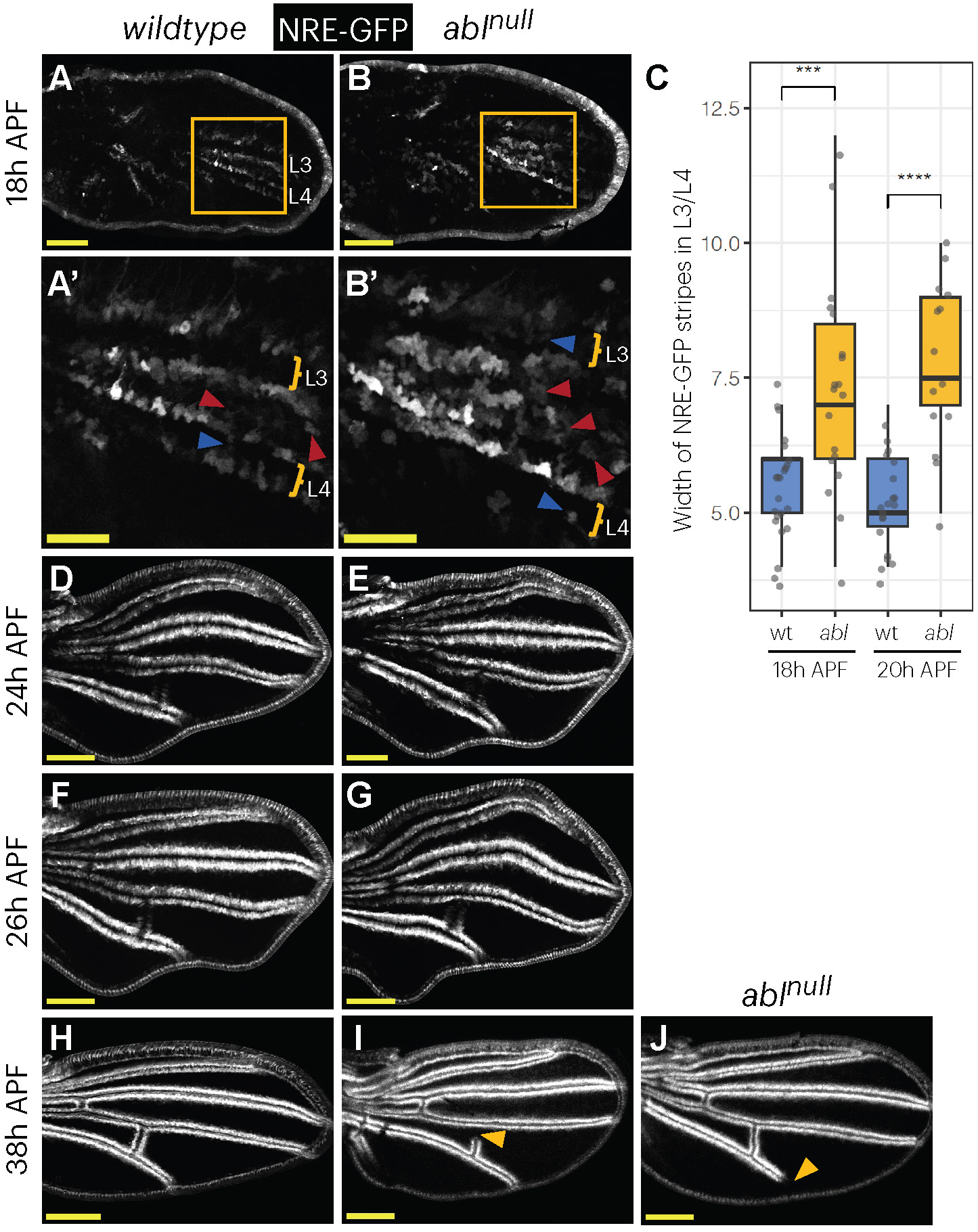
Abl is required to limit Notch signaling during wing vein patterning. (A,B,D-J) Maximal projections of wildtype and *abl^null^* pupal wings. (A,B) 18h APF, with zoomed insets in (A’,B’). (D-E) 24h APF. (F-G) 26h APF. (H-K) 38h APF. Blue and red arrowheads mark ectopic signaling events at intervein and vein cells, respectively. Yellow arrowheads mark a loss of signaling event. Scale bars = 100µm. (C) Plot showing the maximum number of cell rows expressing NRE-GFP reporter in L3/L4 veins at 18h and 20h APF. Each dot represents the measurement from an individual wing. Sample size (# of wings evaluated): wt 18h (N=23), *abl* 18h (N=19), wt 20h (N=20) and *abl* 20h (N=14). A t-student means comparison test was performed and p-values <0.05 (*), <0.01 (**), <0.001(***) and <0.0001 (****) are indicated.

**Figure S2.**
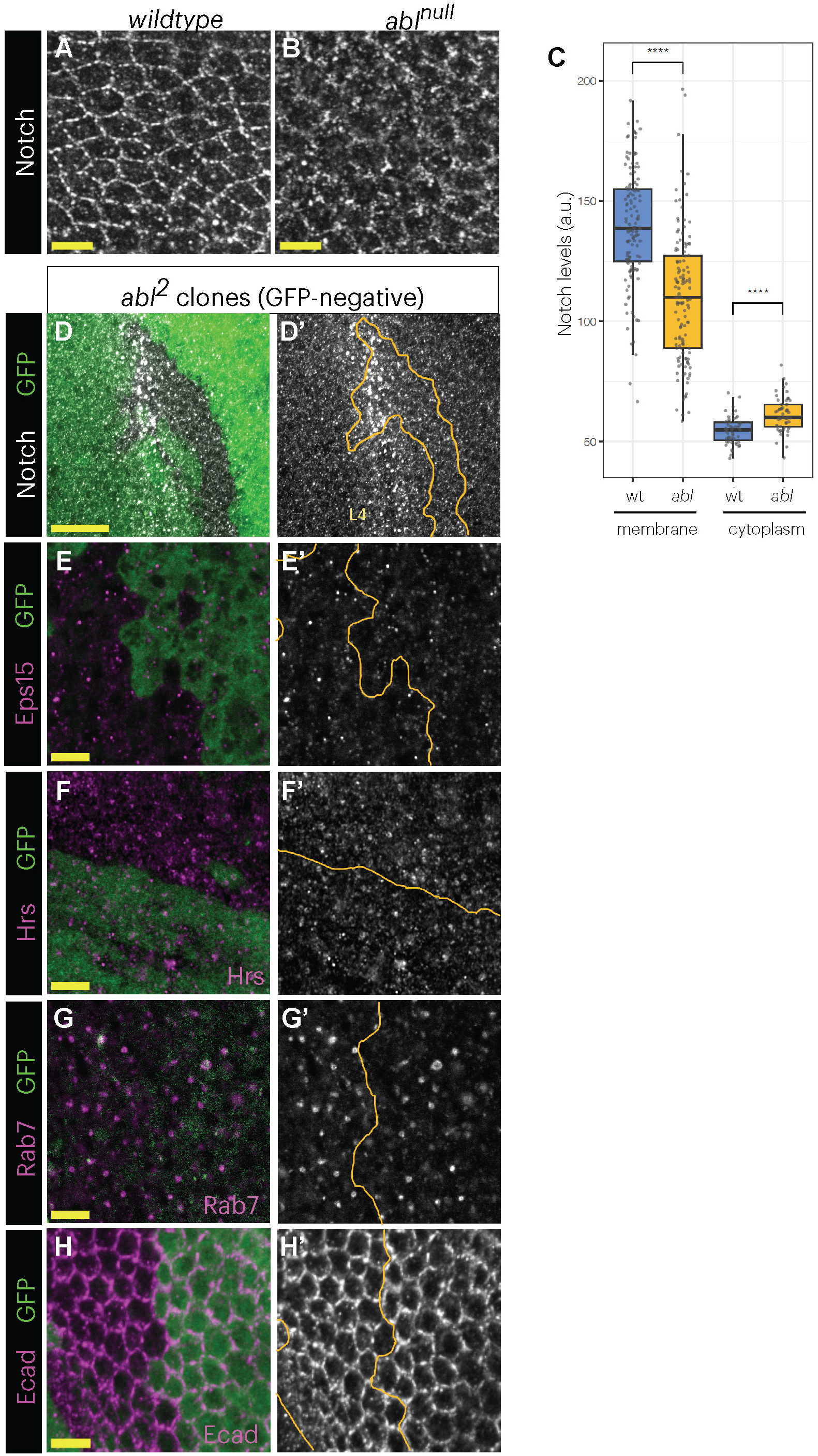
Loss of *abl* selectively perturbs Notch endocytic trafficking (A-B) Maximal projections of wildtype and *abl^null^* pupal wing tissue at 32h APF. Panels shown anti-Notch immunostaining in intervein cells between L3 and L4, proximal to the PCV. Scale bars = 5µm. (C) Quantification of Notch fluorescent intensity at the cortex and cytoplasm of cells shown in (A) and (B). Each dot represents the individual measurement of a cell border (N=130 in wt and N=129 in *abl*) or a cell cytoplasm (N=52 in wt and N=53 in *abl*). A t-student means comparison test was performed and p-values <0.05 (*), <0.01 (**), <0.001(***) and <0.0001 (****) are indicated. Results are similar to those measured at vein areas (Fig. 2E). (D-H) 32hr APF pupal wing tissue with *abl*^2^ null clones marked by lack of GFP (green) and stained with the indicated marker (magenta or white). Yellow lines in D’-H’ mark the clonal boundaries. (D) Notch, (E), Eps15, (F) Hrs, (G) Rab7, (H) E-cad. A maximal projection is shown in (D) and representative single confocal sections are shown in (E-H). Scale bars = 20µm (D) or 5µm (E-H).

**Figure S3.**
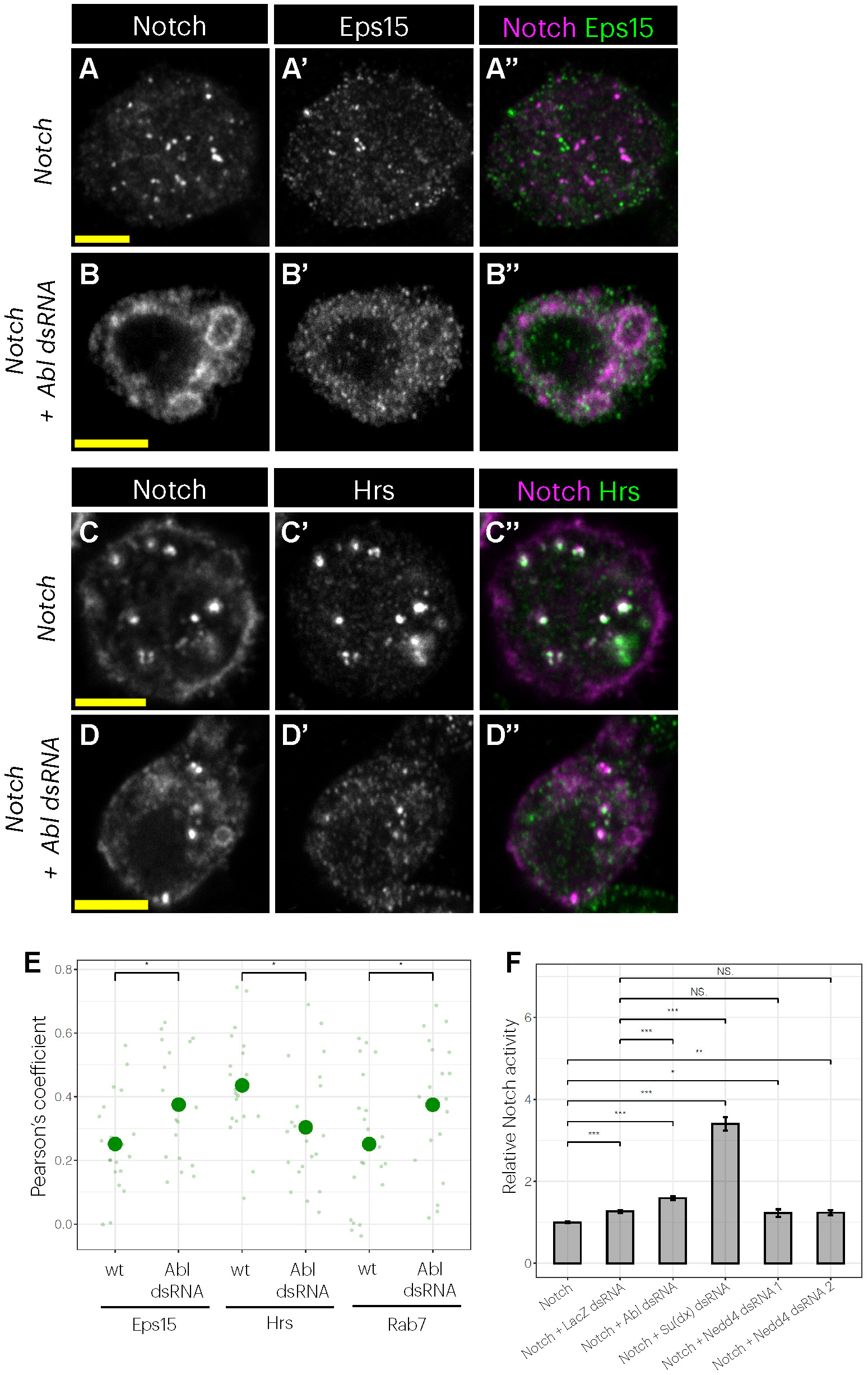
Abl knockdown alters Notch endocytic distribution in S2 cells similarly to *abl* loss in wings. (A-D) Co-imaging of Notch and endosomal markers in wildtype and Abl dsRNA-treated S2 cells. Representative single confocal sections are shown. Transfections: Notch in (A) and (C), and Notch + Abl dsRNA in (B) and (D). Stainings: Notch and Eps15 in (A-B), and Notch and Hrs in (C-D). Scale bars = 5µm. (E) Pearson’s coefficient-based co-localization analysis of Notch and endosomal markers shown in (A-D). Each dot represents the Pearson’s coefficient from an individual S2 cell, with an average of 20 evaluated per condition. A t-student means comparison test was performed and p-values <0.05 (*), <0.01 (**), <0.001(***) and <0.0001 (****) are indicated. (F) Relative Notch activity (NRE>luciferase reporter) produced by co-transfection of dsRNAs targeting Notch trafficking regulators, in comparison to co-transfection of LacZ dsRNA or transfection of Notch alone. A t-student means comparison test was performed and p-values <0.05 (*), <0.01 (**) and <0.001(***) are indicated.

**Figure S4.**
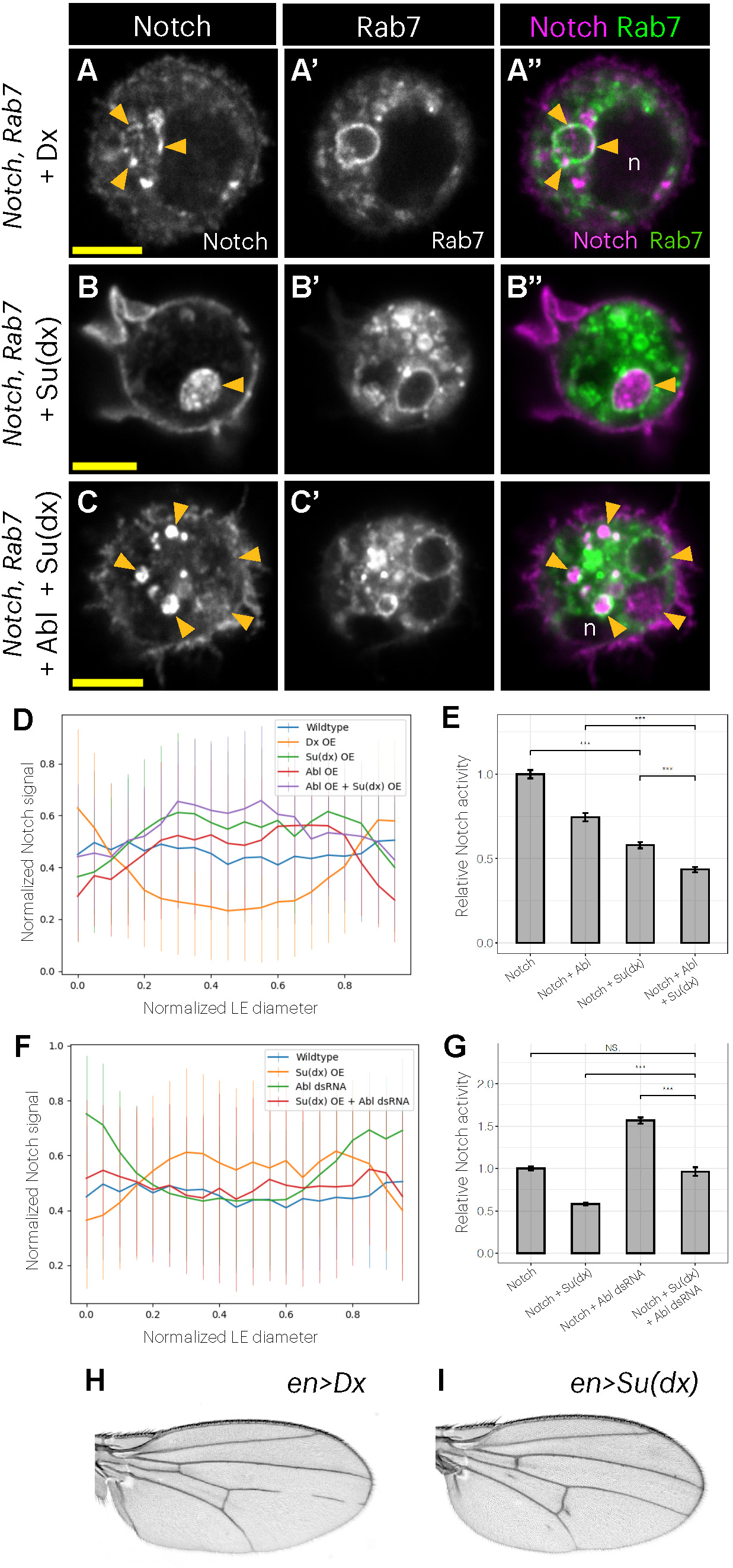
Abl and Su(dx) cooperate to promote Notch internalization into the late endosomal lumen. (A-C) Single confocal sections of representative S2 cells showing Notch (indirect immunofluorescence) and Rab7-EYFP . “n” marks the cell nucleus. Transfections shown: (A) Notch + Dx, (B) Notch + Su(dx), and (C) Notch + Abl + Su(dx). Scale bars = 5µm. (D,F) Plot showing profiles of Notch levels across late endosome diameters in the indicated manipulations. Sample size (# of LEs evaluated): Notch (N=38), Notch + Dx (N=68), Notch + Su(dx) (N=51), Notch + Abl (N=27), Notch + Abl + Su(dx) (N=25), Notch + Su(dx) + Abl dsRNA (N=47). (E,G) Relative Notch activity in the indicated manipulations. A t-student means comparison test was performed and pvalues <0.05 (*), <0.01 (**) and <0.001(***) are indicated. (H-I) en>Dx and en>Su(dx) adult wings, respectively.

**Figure S5.**
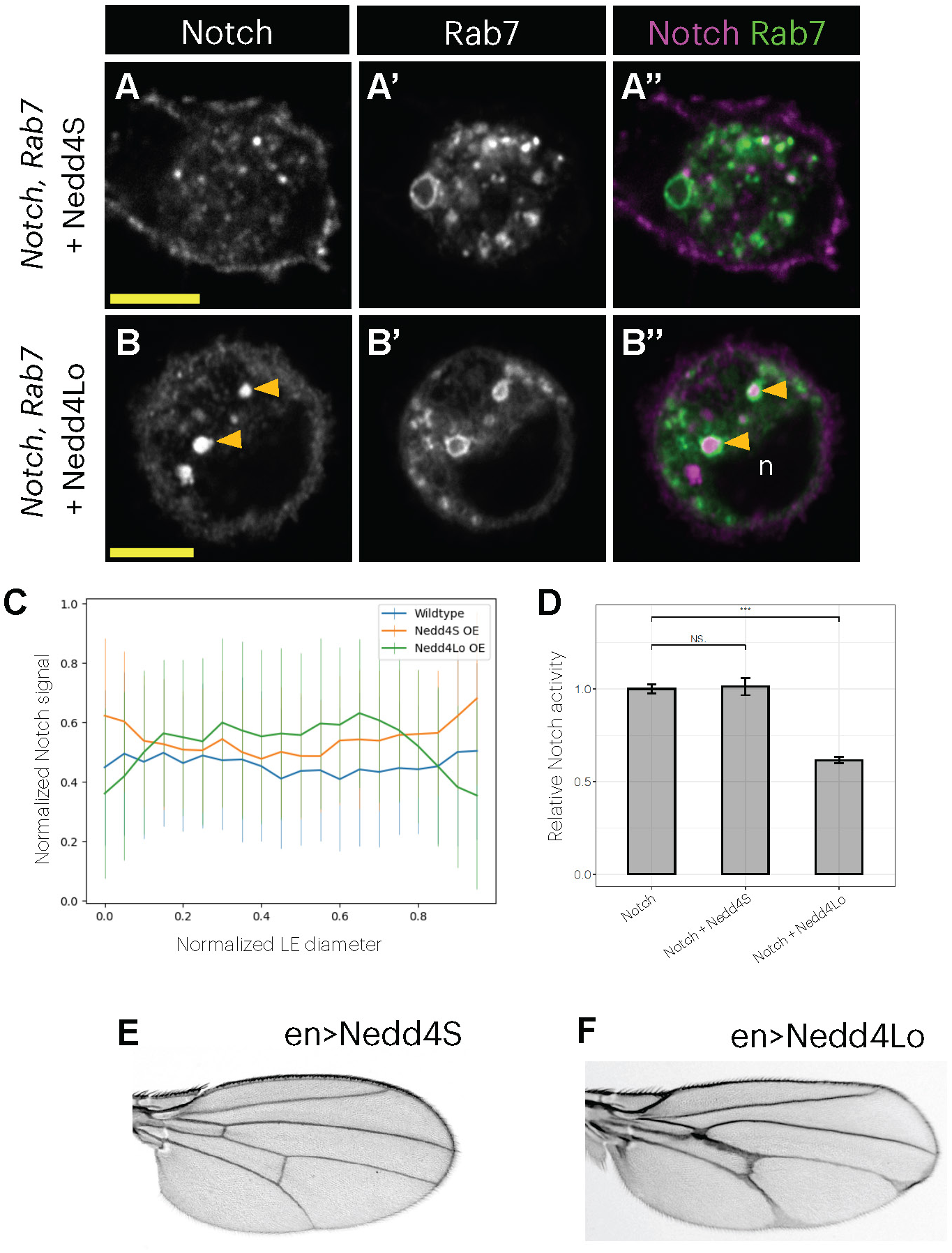
Nedd4Lo promotes Notch internalization into the late endosome lumen to attenuate signaling. (A-B) Single confocal sections of representative S2 cells showing Notch (indirect immunofluorescence) and Rab7-EYFP . “n” marks the cell nucleus. Transfections shown: (A) Notch + Nedd4S, (B) Notch + Nedd4Lo. Scale bars = 5µm. (C) Plot showing profiles of Notch levels across late endosome diameters in (A-B). Sample size (# of LEs evaluated): Notch (N=38), Notch + Nedd4S (N=27), Notch + Nedd4Lo (N=36). (D). Relative Notch activity in manipulations in (A-B). A t-student means comparison test was performed and p-values <0.05 (*), <0.01 (**) and <0.001(***) are indicated. (E-F) en>Nedd4S and en>Nedd4Lo adult wings, respectively.

