## Supplemental Figures 1-5 for "The Abelson kinase and the Nedd4-family E3 ligases co-regulate Notch trafficking to limit signaling"

### Figure S1

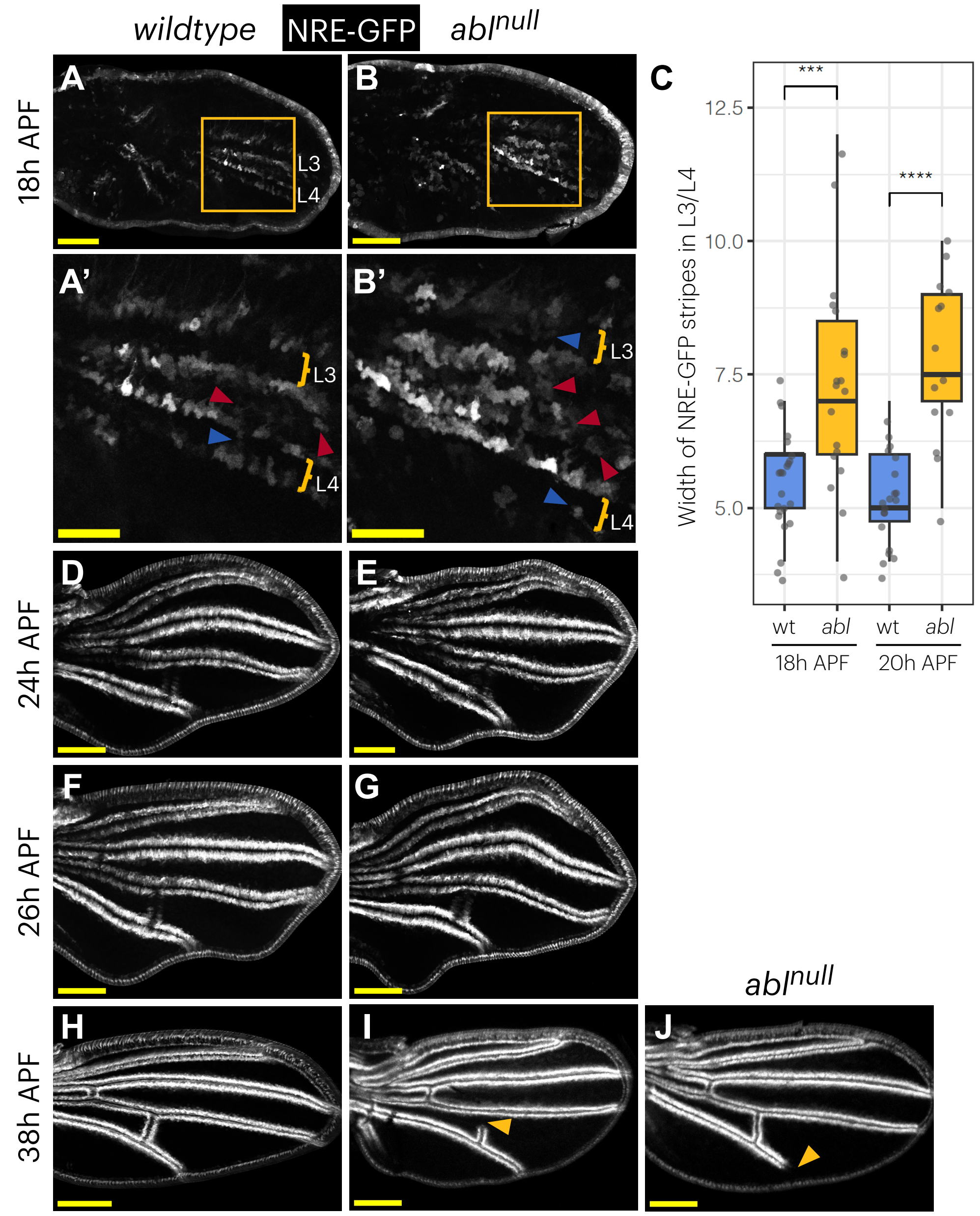

wildtype
NRE-GFP
abl$^{null}$
18h APF
24h APF
26h APF
38h APF
A
B
A'
B'
C
D
E
F
G
H
I
J
L3
L4
Width of NRE-GFP stripes in L3/L4
12.5
10.0
7.5
5.0
***
****
wt
abl
18h APF
20h APF
abl$^{null}$

### Figure S2

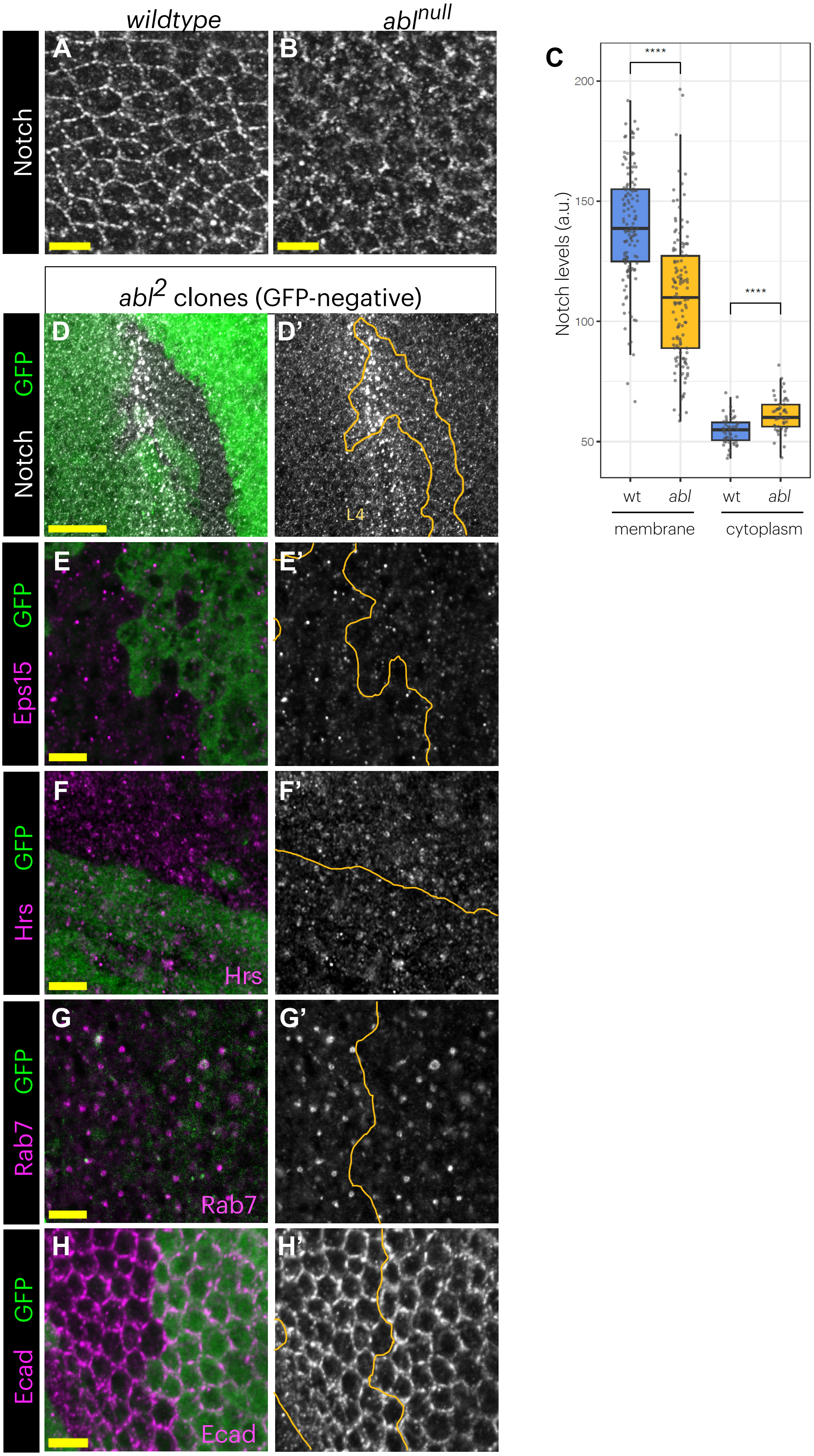

wildtype
abl^null
Notch
A
B
C
****
200
150
100
50
Notch levels (a.u.)
wt
abl
wt
abl
membrane
cytoplasm
abl^2 clones (GFP-negative)
GFP
Notch
D
D'
L4
GFP
Eps15
E
E'
GFP
Hrs
F
F'
Hrs
GFP
Rab7
G
G'
Rab7
GFP
Ecad
H
H'
Ecad

### Figure S3

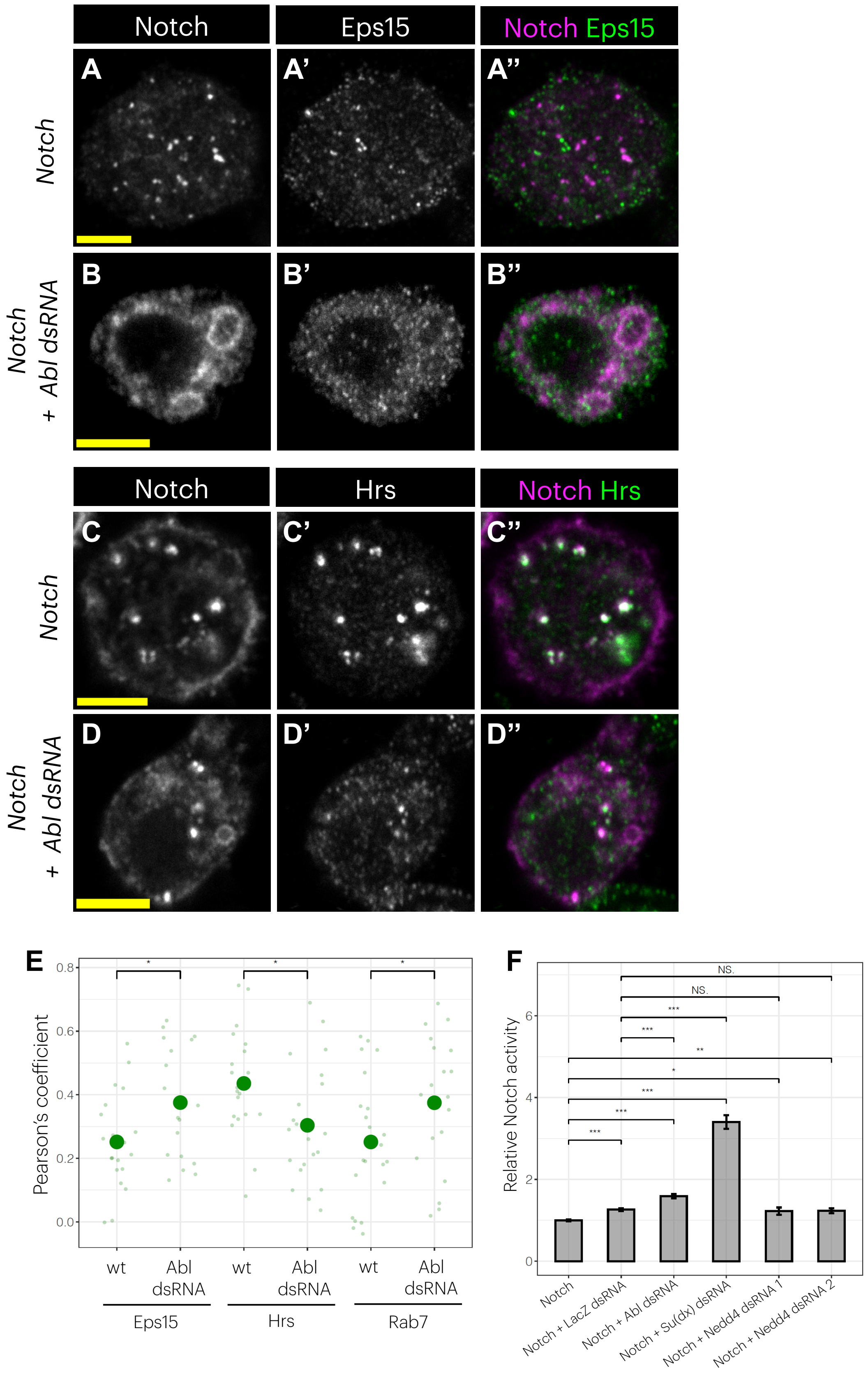
Notch
Eps15
Notch Eps15
A
A'
A''
Notch
B
B'
B''
Notch + Abl dsRNA
Notch
Hrs
Notch Hrs
C
C'
C''
Notch
D
D'
D''
Notch + Abl dsRNA
E
Pearson's coefficient
0.0
0.2
0.4
0.6
0.8
wt
Abl dsRNA
wt
Abl dsRNA
wt
Abl dsRNA
Eps15
Hrs
Rab7
F
Relative Notch activity
0
2
4
6
NS.
NS.
Notch
Notch + LacZ dsRNA
Notch + Abl dsRNA
Notch + Su(dx) dsRNA
Notch + Nedd4 dsRNA 1
Notch + Nedd4 dsRNA 2

### Figure S4

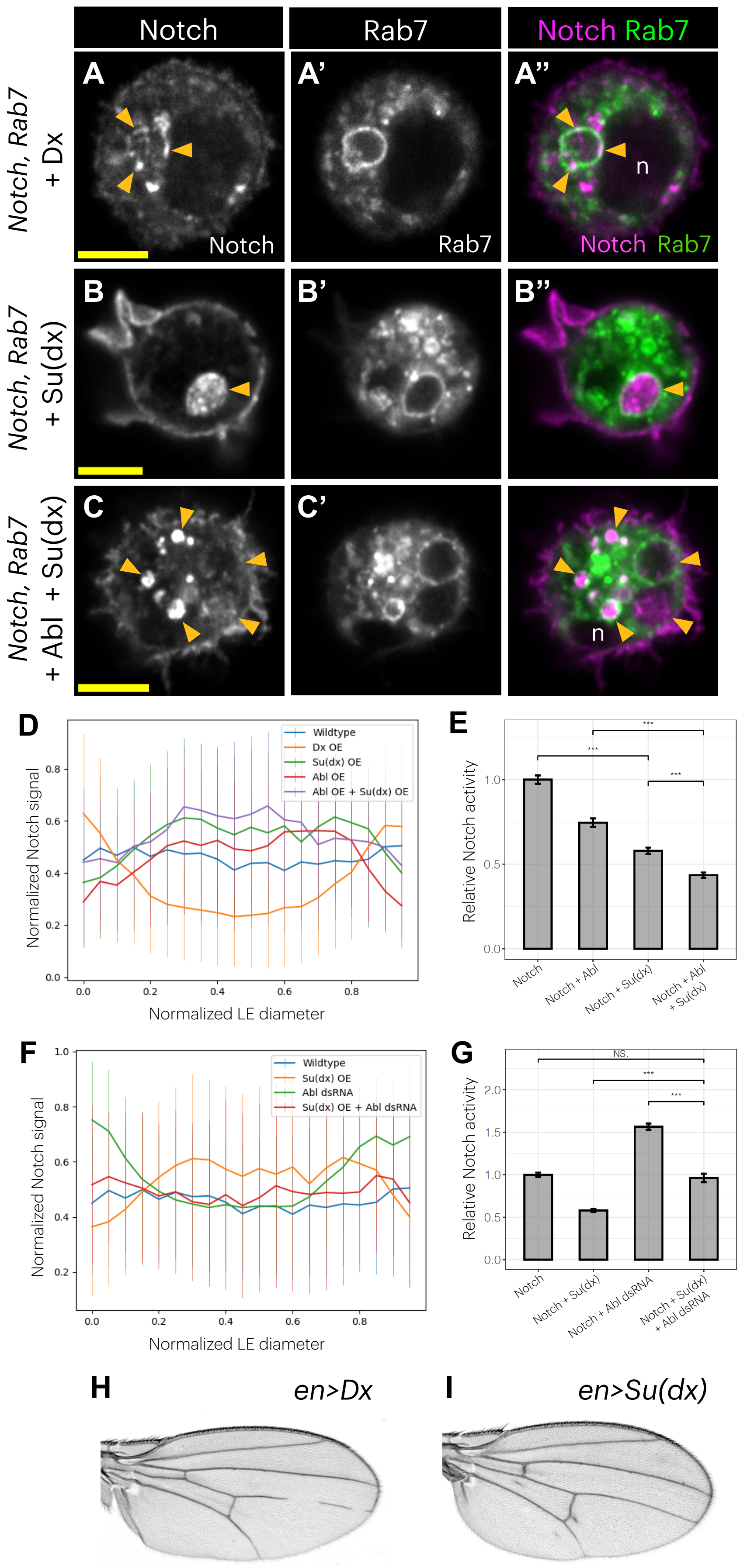
Notch
Rab7
Notch Rab7
Notch, Rab7 + Dx
A
A'
A''
Notch
Rab7
n
Notch Rab7
Notch, Rab7 + Su(dx)
B
B'
B''
Notch, Rab7 + Abl + Su(dx)
C
C'
n
D
Normalized Notch signal
Normalized LE diameter
Wildtype
Dx OE
Su(dx) OE
Abl OE
Abl OE + Su(dx) OE
E
Relative Notch activity
Notch
Notch + Abl
Notch + Su(dx)
Notch + Abl + Su(dx)
F
Normalized Notch signal
Normalized LE diameter
Wildtype
Su(dx) OE
Abl dsRNA
Su(dx) OE + Abl dsRNA
G
Relative Notch activity
NS
Notch
Notch + Su(dx)
Notch + Abl dsRNA
Notch + Su(dx) + Abl dsRNA
H
en>Dx
I
en>Su(dx)

### Figure S5

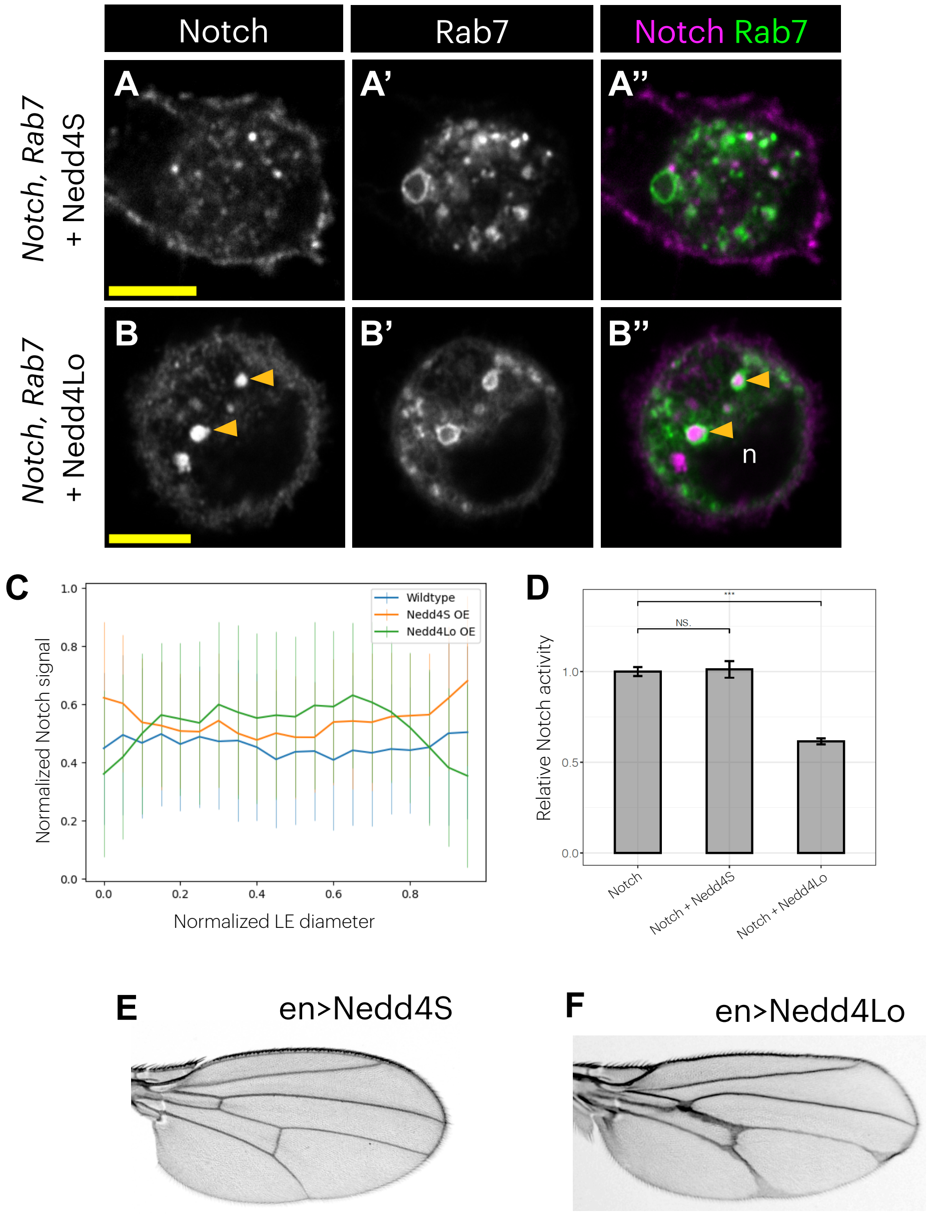
Notch
Rab7
Notch Rab7
A
A'
A''
Notch, Rab7 + Nedd4S
B
B'
B''
n
Notch, Rab7 + Nedd4Lo
C
Wildtype
Nedd4S OE
Nedd4Lo OE
Normalized Notch signal
Normalized LE diameter
D
NS.
***
Relative Notch activity
Notch
Notch + Nedd4S
Notch + Nedd4Lo
E
en>Nedd4S
F
en>Nedd4Lo

**Supplementary Figure Legends**

Figure S1. Abl is required to limit Notch signaling during wing vein patterning.
(A,B,D-J) Maximal projections of wildtype and *abl*$^{null}$ pupal wings. (A,B) 18h APF, with zoomed insets in (A',B'). (D-E) 24h APF. (F-G) 26h APF. (H-K) 38h APF. Blue and red arrowheads mark ectopic signaling events at intervein and vein cells, respectively. Yellow arrowheads mark a loss of signaling event. Scale bars = 100μm.
(C) Plot showing the maximum number of cell rows expressing NRE-GFP reporter in L3/L4 veins at 18h and 20h APF. Each dot represents the measurement from an individual wing. Sample size (# of wings evaluated): wt 18h (N=23), *abl* 18h (N=19), wt 20h (N=20) and *abl* 20h (N=14). A t-student means comparison test was performed and p-values <0.05 (*), <0.01 (**), <0.001(***) and <0.0001 (****) are indicated.
